# HP1β and H3K9me3 Regulate Olfactory Receptor Choice and Transcriptional Identity

**DOI:** 10.1101/2025.08.21.671605

**Authors:** Martín Escamilla-del-Arenal, Rachel Duffié, Hani Shayya, Valentina Loconte, Axel Ekman, Lena Annika Street, Adan Horta, Daniele Canzio, Kevin Monahan, Carolyn Larabell, Marko Jovanovic, Stavros Lomvardas

## Abstract

Diverse epigenetic regulatory mechanisms ensure and regulate cellular diversity. Among others, the histone 3 lysine 9 me3 (H3K9me3) post translational modification participates in silencing lineage-inappropriate genes. H3K9me3 restricts access of transcription factors and other regulatory proteins to cell-fate controlled genes. In mice, olfactory sensory neurons (OSN) express one olfactory receptor (OR) gene out of 2,600 possibilities. This monoallelic and stochastic OR choice happens as OSNs differentiate and undergo dramatic changes in nuclear architecture. OR genes from different chromosomes converge into specialized nuclear bodies and chromatin compartments as H3K9me3 and chromatin binding proteins including heterochromatin protein 1 (HP1) are incorporated. In this work, we have uncovered an unexpected role for HP1β in OR choice and neuronal identity that cannot be rescued by HP1α *in vivo*. With the use of a conditional knock-in mouse model that replaces HP1β for HP1α, we observe changes in H3K9me3 levels, DNA accessibility, and Hi-C contacts over OR gene clusters. These changes alter the expression patterns that partition the mouse olfactory epithelium into five OR expression zones, which results in a reduced OR repertoire leading to a loss of olfactory sensory neuron diversity. We propose that HP1β modulates the competition of OR-promoters for enhancers to promote receptor diversity, by establishing repression gradients in a zonal fashion.

## Introduction

Distinct mechanisms have evolved to generate cell diversity in the animal kingdom. In mammals, two remarkable examples are VDJ recombination in the immune system and OR choice in the olfactory system, where diversity is achieved by different processes (reviewed in 1). Monoallelic and stochastic gene choice are essential for dosage compensation, cell differentiation and cell identity (2–9). Decoding the mechanisms that govern stochastic choice will not only clarify this fascinating mode of gene expression regulation but also reveal fundamental principles of eukaryotic gene regulation.

Stochastic choice is important for neuronal diversity (4,6–8). Singular OR choice is essential for proper function of the olfactory system, as it determines the ligand(s) from the environment to which the OSN will respond and ensures axonal wiring to a stereotyped glomerulus in the olfactory bulb (10). Expressing multiple ORs per cell could lead to axonal wiring defects and errors in transmitting information (11). While in a given OSN, OR choice is random and stochastic, each OR is restricted in expression to one of five anatomical regions in the Main Olfactory Epithelium (MOE). The physical location within the tissue determines which set of ORs have the potential to be expressed in a given cell. Nuclear organization and heterochromatinization of the OR gene loci are essential for both silencing and enabling the activation of an OR allele (12–14). OR genes form interchromosomal contacts with at least 64 OR enhancers, located all around the genome (15, 16). Transcriptional upregulation of one OR allele serves to silence other OR alleles, possibly through competition with other OR-enhancer complexes (17). At this point, the Unfolded Protein Response signaling pathway (18, 19) locks in the OR selection.

Recent work has revealed zonal differences in H3K9me3 deposition patterns in the MOE (14), raising the possibility that heterochromatin plays a fundamental role in zonal OR choice and neuronal identity. Regulation of cell fate and cell identity is associated with chromatin by promoting or restricting access of DNA-interacting proteins to their target sequences (19). This facilitates the expression of lineage specific genes, in response to signals that are activated during developmental processes (16). Previous work shows that heterochromatin regulates OR choice (20). The transcriptionally silent OR genes are marked by H3K9me2 and acquire H3K9me3 upon differentiation (27). The first insight that H3K9me3-based heterochromatin OR gene silencing was necessary to generate gene diversity, came from G9a and GLP histone methyltransferases (HMT) conditional knock-out (KO) alleles (20). In the absence of G9a and GLP, which are necessary for H3K9me2 incorporation, H3K9me3 is not established in OR gene clusters. The absence of H3K9me3 interfered with normal monogenic receptor expression and led to a decrease in OSN diversity (20). However, the mechanism by which H3K9me3 impacts OR gene choice remains unknown.

The HP1 proteins are highly conserved nonhistone chromosomal protein, that bind H3K9me2/3 and are involved in heterochromatin-mediated gene silencing (21–24). Mammalian cells contain three closely related HP1 homologs, HP1α (Cbx5), HP1β (Cbx1), and HP1γ (Cbx3) (23). These HP1 isoforms share two highly conserved domains, an N-terminal chromo-domain and a C-terminal chromo shadow domain, separated by a hinge region. The chromo-domain is responsible for H3K9me2/3 binding (25, 26), while the chromo shadow domain is used to form HP1 dimers and as a platform to recruit HP1- binding partners, including transcriptional co-repressors and H3K9 HMT (21, 27–29). However, HP1 protein function in establishing cell diversity and cell identity is still not fully understood.

In the current study we demonstrate that HP1β facilitates the incorporation of H3K9me3 methylation in neurons from the MOE, regulating OR choice and transcriptional identity. The deletion of the murine *Cbx1* gene, which encodes the HP1β protein, results in perinatal lethality (30). To overcome this problem, we designed a knock-in mouse model that after CRE expression rescues the lethality observed in the HP1β KO mouse, with the expression of HP1α. HP1α and HP1β proteins were localized in different nuclear compartments, highlighting the compartmentalization of these two proteins and their function. In the absence of HP1β, we observe a drop in H3K9me3 at OR clusters as well as a drop in genome-wide heterochromatin levels, impacting nuclear architecture, OR gene choice and transcriptional identity of the cell. We propose a model for HP1β in regulating OR choice and OSN transcriptional identity.

## Results

### Developmental switch from HP1α and HP1β proteins during OSN differentiation

Terminally differentiated cells require H3K9me3-mediated heterochromatin silencing for lineage commitment and maintenance of cell identity (31). As OR alleles are chosen for expression in immature OSNs, the alleles that are not selected in a given neuron will be enriched for H3K9me3 and recruited to a compact heterochromatic focus (11, 19). Given the co-localization of HP1β to these chromatin foci, we hypothesize a function of this family of proteins in OR regulation. RNA-Seq levels of *Cbx1* (gene encoding HP1β) and *Cbx5* (HP1α) from fluorescence activated cell sorted (FACs) populations during OSN differentiation suggest they have a role in two distinct developmental windows (Fig.1A). *Cbx1* RNA expression peaks in the Mash1 (also known as Ascl1) cells and Mash1/Neurogenin-1 double positive cells, coincident with the step where stem cells commit to the neuronal lineage and undergo a dramatic reorganization of heterochromatin (12). *Cbx1* RNA remains abundant in immature (Neurogenin1 positive and Atf5 translating populations) and mature olfactory sensory neurons (OMP positive population). The predominant presence of HP1β in the neuronal layers suggests a prominent role for HP1β and not HP1α in OR regulation, but we do not exclude HP1α from having a role in setting up the system in early stages of differentiation. To visualize HP1α and HP1β localization in the olfactory epithelium, we performed Immunofluorescence (IF) (Fig.1B, Fig.1C) and quantified staining intensity from basal to apical positions in the tissue section (Fig1.D). HP1α staining is prominent throughout the nucleus in basal cells of the olfactory epithelium, where stem cells reside, and in the sustentacular support cells which are most apical in the tissue. With differentiation, HP1α undergoes a dramatic change in localization, moving with the pericentromeric chromatin to the heterochromatic foci (Fig.1C) and gradually fades out from the system as olfactory neurons mature, at the mRNA levels (Fig.1A) and at the protein levels (Fig.1B). On the other hand, HP1β exhibits staining throughout the neuronal layer, and in contrast to HP1α, is excluded from non-neuronal basal cells (Fig.1C). Remarkably, in immature neurons HP1β localizes to the heterochromatic foci and with maturation it is found to encircle the heterochromatic foci of mOSN in the MOE (Fig.1C, Fig.1D, compare HP1β localization to Hp1α in iOSN vs mOSN). This staining pattern is consistent with the localization of the olfactory receptor gene loci, around the DAPI dense heterochromatic focus, as seen by DNA-FISH with a “pan-OR” probe, which recognizes many members of the olfactory receptor gene family (12). Here we showed HP1α and HP1β have distinct developmental windows of action. This result shows a developmental switch in HP1 proteins during differentiation, with distinct developmental windows, except for basal cells, where HP1α and HP1β both mark newly formed chromatin foci (Fig.1C). We propose independent roles for these proteins in OR gene regulation.

**Figure 1.**
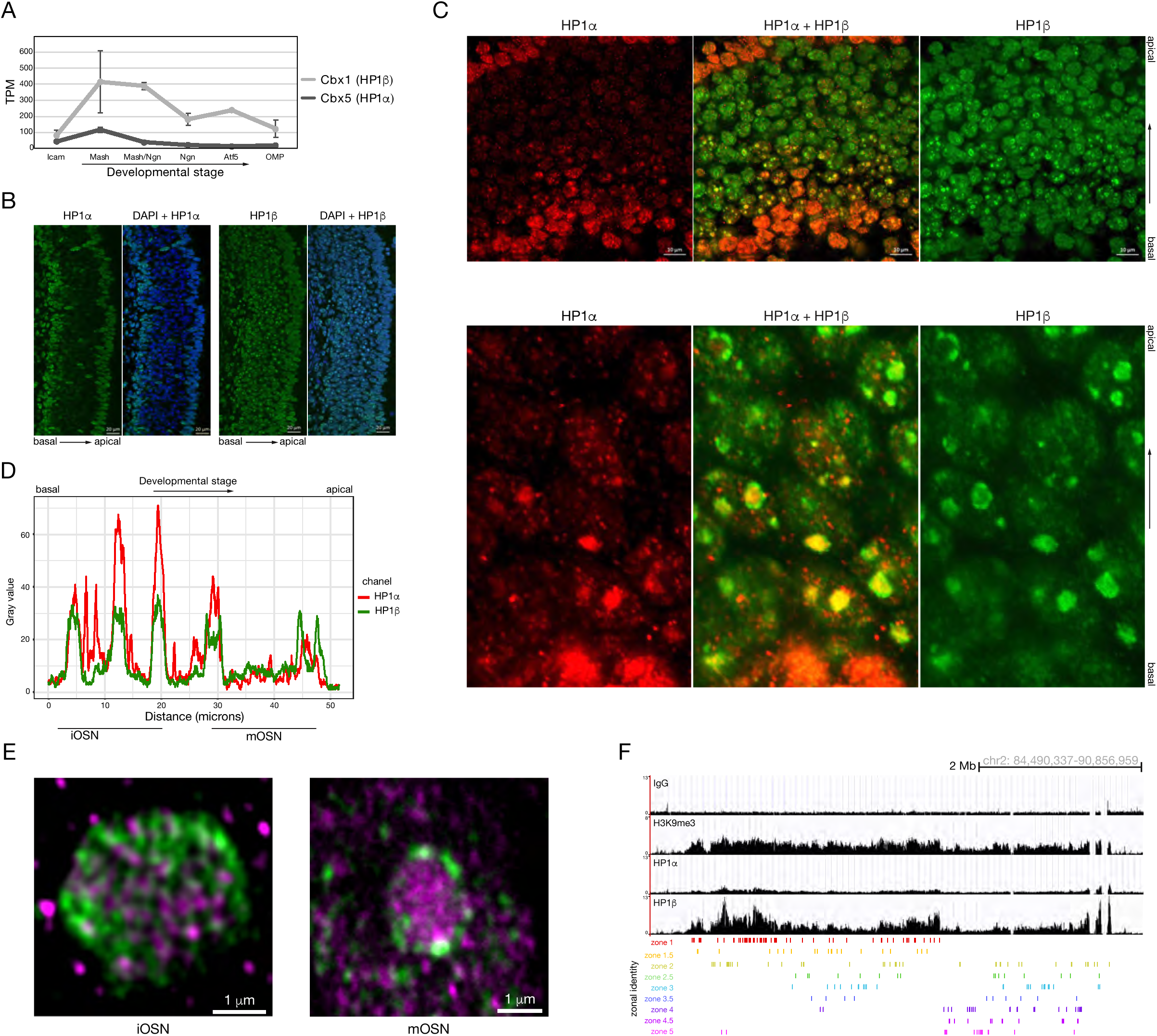
HP1α and HP1β segregate into different compartments during OSN differentiation. A) RNA-seq of sorted cells at different points in olfactory sensory neuron differentiation. TPM (Transcript Per Million). Data from two biological replicates. B) IF on Mouse Olfactory Epithelium (MOE) sections, using specific antibodies against HP1α and HP1β. Basal and apical localization of the tissue is indicated. Scale bar 20μm. C) IF on MOE sections using specific antibodies against HP1α (red) and HP1β (green). Top images taken with a 40X lens, shows HP1α and HP1β staining colocalize in immature Olfactory Neurons (iOSN). Scale bar 10μm. Bottom images taken with a 64X lens, shows HP1α and HP1β staining is dynamic, with HP1β moving to the periphery, localizing these two proteins to distinct nuclear compartments that resembles models of phase transition and suggests a role in 3D organization. D) IF staining intensity quantified from bottom image on “C” shows HP1α and HP1β colocalization kinetics. E) SoRA Spinning disk high resolution microscopy from an iOSN focus (left) and a mOSN (right). Scale bar 1μm. F) ChIP-seq signal tracks from chromosome 2 OR cluster. Values are reads per 10 million. Below the signal tracks, OR genes are depicted in different colors indicating the assigned expression zone. Antibodies used on IF: HP1α (ab109028) and HP1β (ab10811). Antibodies used on ChIP: H3K9me3 (Ab8898) HP1α (ab109028) and HP1β (D2F2, Cell Signaling).

### HP1α and HP1β localize to different nuclear compartments

Whether or not HP1α and HP1β have redundant roles *in vivo* has not been described. *In vitro* studies show HP1α can phase transition, however HP1β cannot (32, 33). During the process of differentiation, these two proteins seem to co-localize only in newly formed heterochromatic foci (Fig.1C), opening the possibility for redundant overlapping functions. To gain further insight into the dynamics of these two proteins in the cell nucleus, we analyzed their spatial localization over the course of OSN differentiation. Using super resolution confocal microscopy, we examine HP1α and HP1β sub-nuclear localization. Remarkably, at higher resolution in iOSNs we observe complete segregation of these two proteins in the heterochromatic foci (Fig.1E and SFig.1B), suggesting that HP1α and HP1β mark different types of heterochromatin compartments, and pointing for independent roles in OR regulation. This parsing of HP1α and HP1β into different compartments in a single chromatin focus, followed by the migration of HP1β to the periphery of the chromatin focus in mature OSNs (Fig.1E and SFig.1C), led us to consider HP1β’s role in nuclear architecture and foci formation (see later). The fact that the OR clusters migrate to the periphery in parallel with HP1β and that HP1α does not share compartment localization with HP1β suggests a prominent role for HP1β in OR regulation and not HP1α. This would mean that HP1α does not interact with the OR gene clusters. To test this hypothesis, we performed ChIP-seq from isolated pure populations of differentiating cells using FACs (Fig1.F, SFig.2). We observed high levels of HP1β protein incorporated into OR gene clusters compared to HP1α. Taken together, these results suggest that HP1 proteins have different functions in OR regulation with a major role for HP1β.

**Figure 2.**
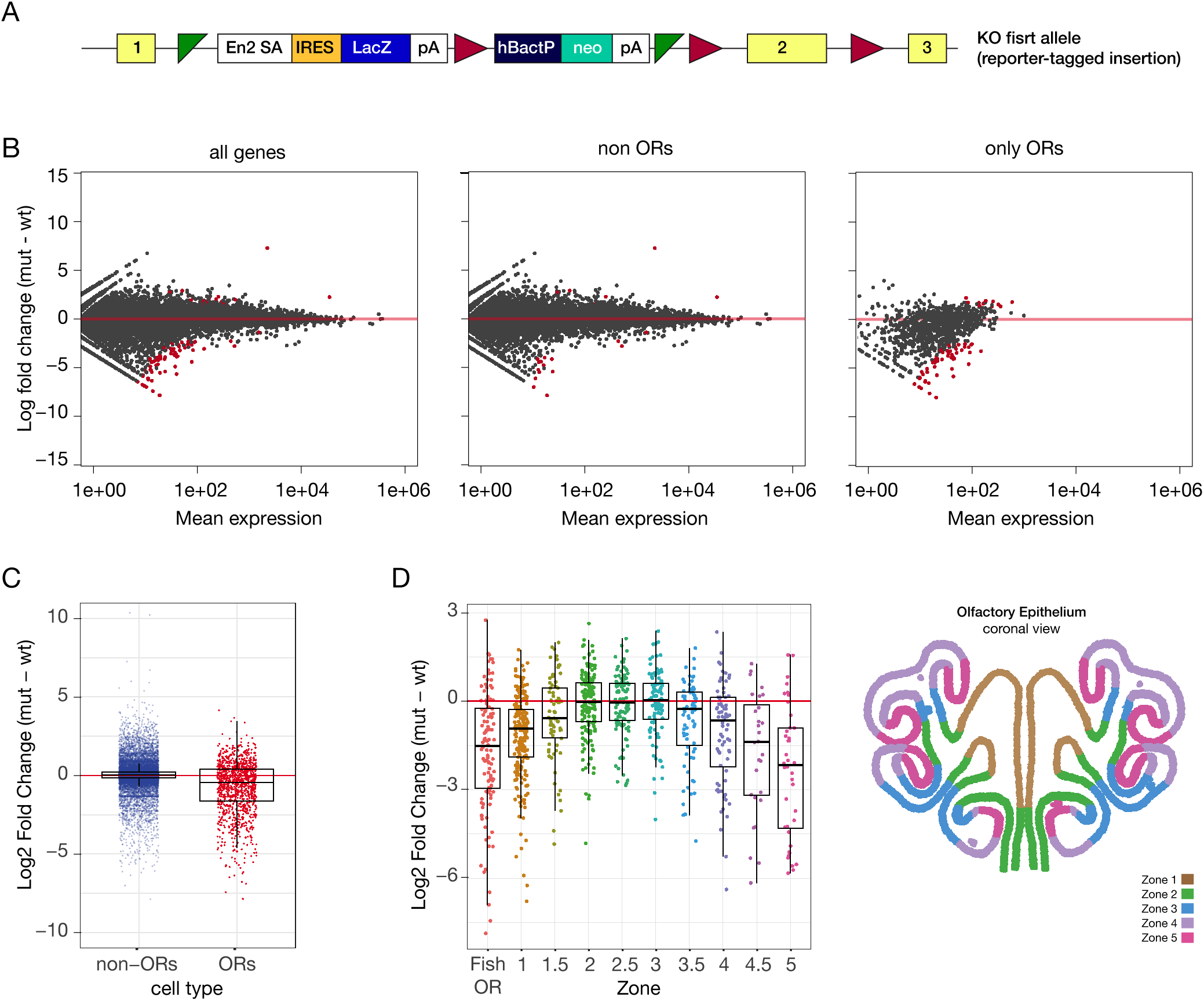
OR expression is affected in a HP1β KO mouse model. A) Diagram of HP1β KO allele (from Sanger) used to study the dependence of olfactory choice regulation on HP1β. B) RNA-seq analysis of gene expression in HP1β KO versus littermate controls. Bulk RNA-seq on E17.5 embryos. Significantly changed genes are colored red (Padj < 0.05 for greater than 1.5-fold change, Wald test, n = 3 biological replicates. C) Scatter plot of bulk RNA-seq on E17.5 embryos comparing HP1β KO versus littermate controls. D) RNA-seq reads were bioinformatically separated into zones to which the OR receptors belong. Right, a diagram depicting the MOE divided into transcriptional zones is shown for the readers’ reference.

### Constitutive depletion of HP1β during embryogenesis leads to a zonal effect on OR expression

A previous study reported that HP1α constitutive knockout (KO) mice are viable with no obvious phenotype, while HP1β constitutive KO mice die at birth or a few hours after (30). This lethality could be due to anosmia, which would prevent pups from suckling, but the lethality can also be associated with genome instability (30). To test a possible role from HP1β in OR regulation, we turned to a knockout-first allele HP1β mouse from Sanger (Fig.2A and Supplemental material), which exhibited embryonic lethality. We dissected the olfactory epithelium from embryonic day (E)17.5 *Cbx1* KO embryos and their wildtype littermates. RNA-Seq data from whole olfactory epithelium showed a specific reduction of 4.3% in OR expression in the *Cbx1* KO E17.5 embryos compared to control littermates (Fig 2B), with minimal effect of 0.07% change in non-OR gene expression (Fig.2B and Fig.2C). This result supports HP1β’s role in olfactory receptor regulation, while raising questions about how loss of a repressive heterochromatin factor would reduce gene expression. Systems where HP1β seem to work as a trans-activator have been reported (4). To further parse OR dysregulation in the *Cbx1* constitutive KO, the olfactory receptor genes were bioinformatically divided into anatomical zones, based on published data (35 and Fig.2D). Class I ORs, the most evolutionarily ancient members of this gene family, and zone-1 ORs exhibited reduced expression in the *Cbx1* KO, whereas zone-2 and zone-3 ORs were largely unaffected. Zone-4 and zone-5 ORs were expressed at greatly reduced levels compared to control littermates (Fig.2D). We also examined differentiation makers including Ngn-1 and OMP and found no significant effect on these differentiation markers (not shown). These data reveal a zonal effect on OR gene expression upon loss of HP1β.

### Zone-4 and zone-5 ORs become activated as *Cbx1* and *Cbx5* levels swap

HP1β constitutive knockout mice die *in utero*, at a stage when olfactory receptor expression is low compared to post-natal embryos (SFig.3A and SFig.3B). At birth, olfactory receptor expression increases up to 5-fold over embryonic levels (SFig.3A and SFig.3B). Interestingly, OR activation is zonal (SFig.3C), with receptors from zone-4 and zone-5 exhibiting higher levels of activation after birth compared to the other anatomical zones (while in embryos zone-1 to zone-3 OR had higher levels of expression compared to zone-4 and zone-5 OR’s). This result shows how activation of the olfactory system does not happen synchronously. The developmental formation of this tissue could explain zonal activation, where the more dorsal parts of the tissue form first. Cbx1 and Cbx5 expression levels swap at birth, with Cbx5 present as with higher levels in E17.5 embryos and Cbx1 becoming the dominant paralog after birth (SFig.3D, SFig.3E). This result is concomitant with the activation of zone-4 and zone-5 ORs (SFig.3D, SFig.3E) as OR differentiation and choice take place.

**Figure 3.**
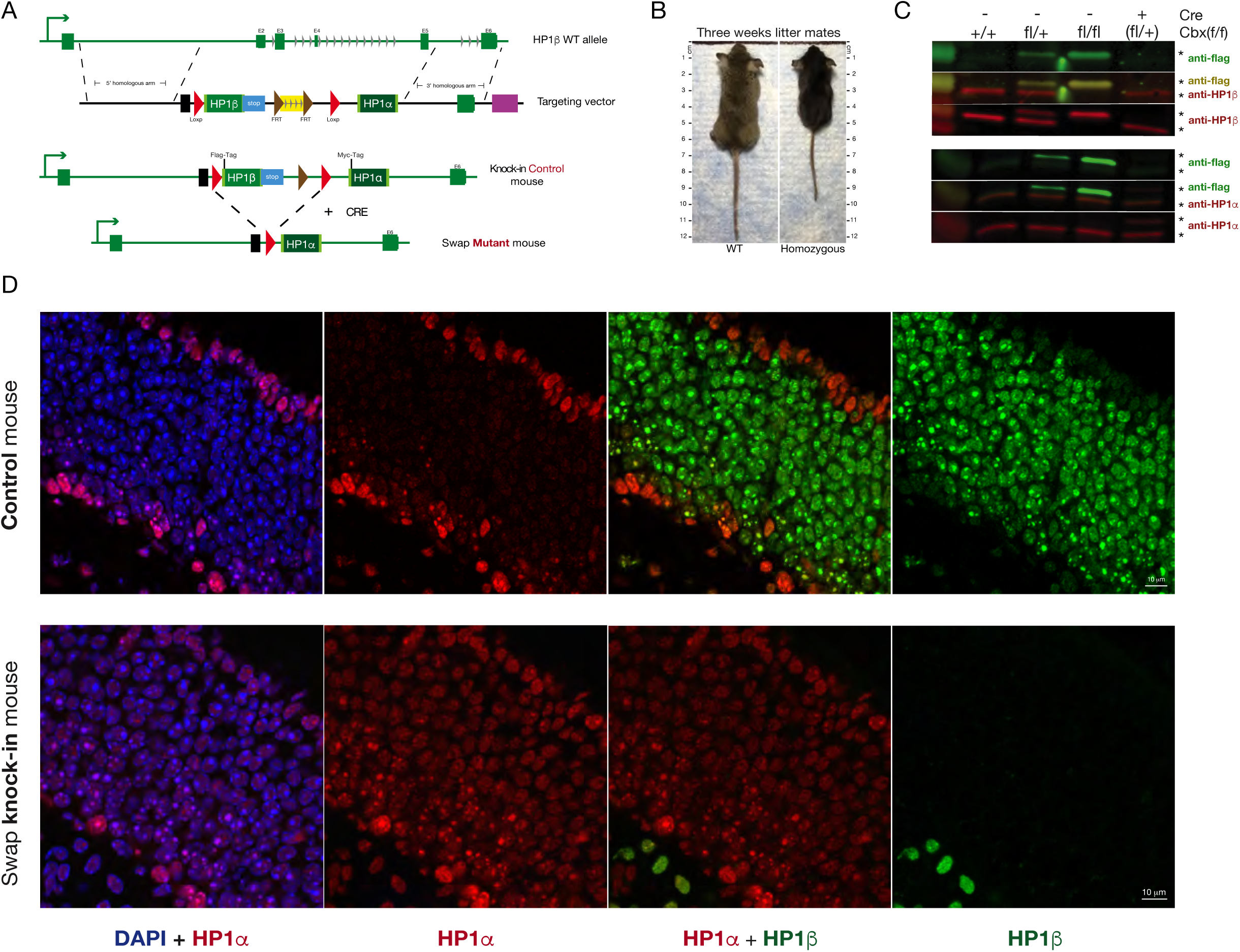
HP1β swap by HP1α results in phenotypic defects and high lethality. A) Schematic representation of the the swap mouse design. We targeted the HP1β locus with a construct containing the HP1β coding sequence, followed by a 3xStop codon and followed by HP1α coding sequence. After CRE recombination, HP1α will replace HP1β and the 3xStop codon removed. Hp1α knock-in will be regulated by HP1β endogenous promoter. B) Picture from three weeks swap vs ctrl mouse littermates, shows a phenotypic effect on the size of the mutant mouse. C) Western Blot on MOE protein extracts from WT (+/+), heterozygous (fl/+), and homozygous mice (fl/fl), with the specified antibodies. D) IF on HP1 knock-in (control) mouse (top panels) or after CRE recombination (bottom panels) using HP1α (red) or HP1β (green) specific antibodies. Antibodies used on WB: Anti-Flag (SIGMA M2 F-3165), HP1α (ab109028) and HP1β (ab10811). Antibodies used on IF: HP1α (ab109028) and HP1β (ab10811).

### HP1α expression partially rescues high lethality observed in the HP1β KO mice

To determine whether HP1α could rescue HP1β induced lethality, and to gain deeper insight into how these proteins regulate OR choice and neuronal identity, we sought to develop a mouse that would substitute HP1β for HP1α, and mimic the spatiotemporal expression pattern of HP1β during OSN differentiation. To this end, we generated a conditional knock-in mouse to simultaneously delete *Cbx1* and express *Cbx5* from the *Cbx1* endogenous promoter (Fig.3A). The inserted construct has the coding sequence of HP1β followed by 3X stop codons and flanked by two flox elements. The HP1α coding sequence is downstream of the second flox element.

In the absence of Cre, the HP1β knock-in is expressed from the endogenous HP1β promoter, and the 3X stop prevents HP1α from being expressed (Fig.3C and Fig.3D upper panel) (referred to as “control mouse” throughout). In order to understand if the knock-in construct has an effect on the RNA expression or protein levels, RNA-seq and Western Blot comparing WT versus the control mouse were done and no significant differences were found (SFig.4A, SFig.4B and Fig.3C). IF analysis shows similar HP1α and HP1β staining compared to WT (compare Fig.3D upper panel, with Fig.1C) as well as similar H3K9me3 staining (data not shown). Western Blot on heterozygous mice containing one copy of *Cbx1* and one copy *Cbx5* Myc tagged protein, showed comparable protein levels between the endogenous and the introduced proteins (Fig.3C, in Cbx^fl/+^ compere staining with anti-flag or an anti-HP1β). Size and behavior between WT and control mice are indistinguishable (data not shown). Taken together these data confirm that the knock-in control mouse is comparable to a WT mouse.

**Figure 4.**
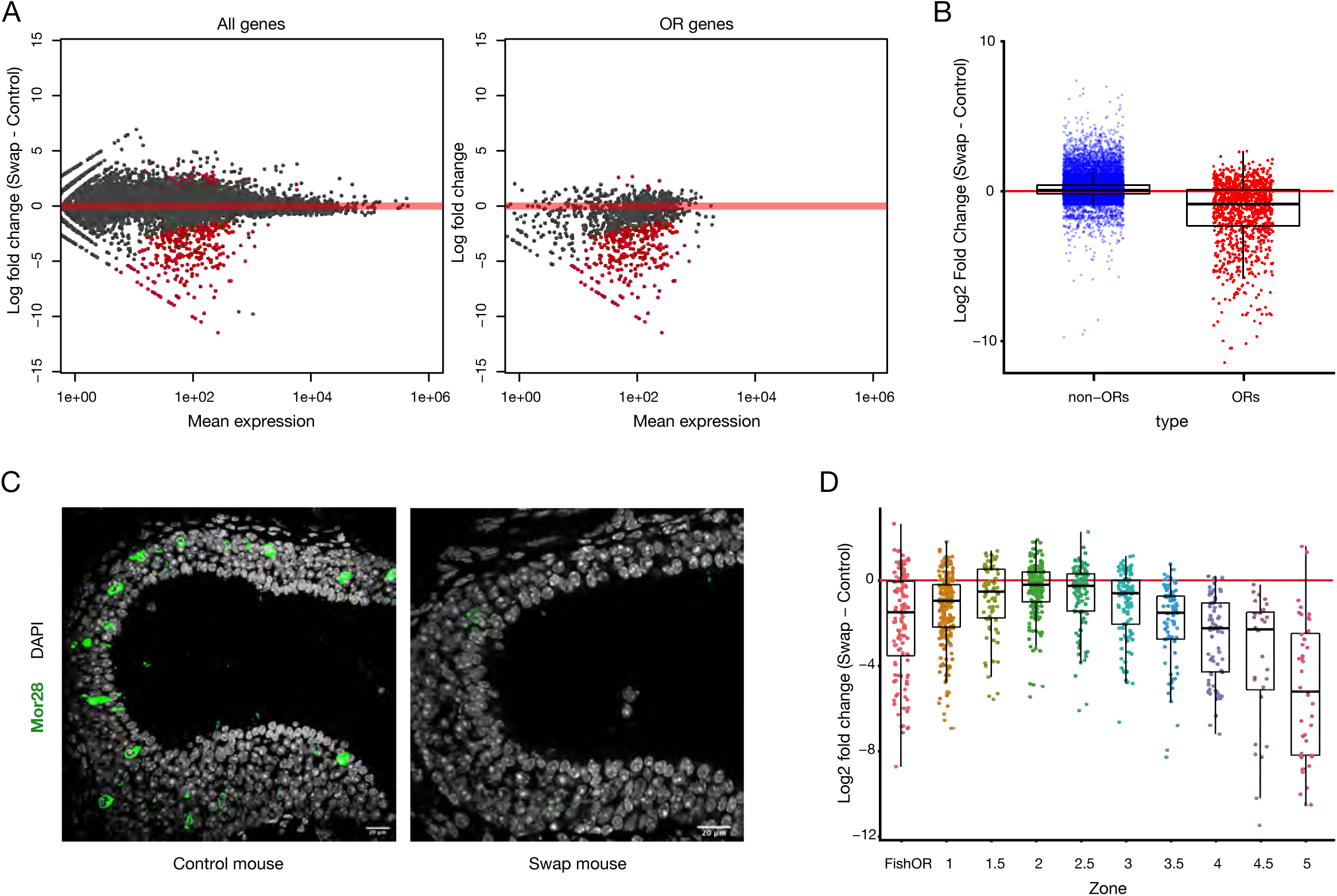
HP1β has a specific role in OR gene regulation that HP1α cannot rescue. A) RNA-seq analysis of gene expression from sorted mOSN cells from swap mouse versus control mouse. Significantly changed genes are colored red (Padj < 0.05 for greater than 1.5-fold change, Wald test, n = 3). B) Boxplot of RNA-seq from swap mouse vs control quantifying non-OR genes and OR genes. C) IF with a zone-5 OR specific antibody (mor28) in zone-5 tissue sections, comparing control versus swap mouse. Scale bar is 20μm. D) Boxplot comparing OR gene expression (Log2FC Swap - Control). RNA-seq reads were bioinformatically separated into the zones to which the OR receptors belong. HP1β KO effects on OR expression is zonal.

The *Foxg1Cre* allele (36) was used to conditionally delete *Cbx1* and “rescue” with *Cbx5*, in neuronal lineages of the mouse, which includes the olfactory sensory neurons. Cre positive progeny with two “HP1 swap” alleles (termed “swap mice” throughout) express HP1α wherever HP1β was expressed and do not express HP1β (Fig.3D, lower vs upper panels). Upon CRE recombination we rescue the lethal phenotype observed in the *Cbx1* KO (Fig.2), however swap mice are runts and do not thrive as well as heterozygote or Cre negative littermate controls. Few swap mice are viable until adult life, and most die between post-natal (PN)0 and PN21. This reduced viability could be an effect of deleting HP1β or by the ectopic expression of the inserted HP1α gene. Heterozygous mice showed no difference in viability compared to WT mice (data not shown), supporting an effect related to the absence of HP1β and not for the introduction of HP1α. IF on swap mice reveals strong staining of HP1α that spreads throughout the neuronal lineage of the olfactory epithelium, in stark contrast to a Cre negative littermate control (Fig.3D and SFig.4B). Importantly, this staining demonstrates that HP1α is now expressed in cells where endogenous HP1β was expressed, however, we do not observe HP1α staining encircling the DAPI dense foci where HP1β is observed. This result suggests differential recruitment of these two proteins to their targets. HP1β staining in the swap mice is absent from the MOE (Fig.3D). As HP1β knockout cannot be fully rescued by HP1α, we suggest HP1β has a specific function in regulating OR expression.

### OR choice is disrupted in the absence of HP1β

To determine whether Hp1α could rescue the effect in OR expression observed in the HP1β KO mice (Fig.2A), we performed bulk RNA-Seq in whole MOE from the swap mice compared to littermate controls. Our data show a dramatic downregulation of 25% of olfactory receptors, with minimal changes to non-OR genes (Fig.4A and Fig.4B). These data further support a role for HP1β in OR regulation that cannot be rescued by HP1α. To confirm this result, we performed IF for the zone-5 OR Mor28 (*Olfr1507*) in zone-5 olfactory epithelium (Fig.4C) and observe a strong reduction in the number of OSNs that choose this receptor compared to littermate controls. The zonal identities of dysregulated ORs were examined, and only zone-2 ORs were expressed at normal levels, comparable with the HP1β constitutive KO (Fig.2D). ORs from other zones showed decreased expression as a class, with zone-5 ORs being the most strongly affected (Fig.4D). These results show a zonal function for HP1β protein in OR regulation, with ORs from ventral zones most strongly dysregulated. To explain this result, we considered different scenarios including cell death, cell maturation, skewed OR choice, or an effect on cell transcriptional identity.

### OR choice and zonal identity are disrupted in the HP1 swap mice

Bulk RNA-Seq data demonstrate a dysregulation of OR genes in the swap mice. Since zone-5 epithelia is present by microscopy analysis, we eliminated cell death as an option and considered three possible explanations for these results: 1) OSNs from zone- 4 and zone-5 are chosen but expressed at a lower level. 2) OR choice is skewed to zone- 1 and zone-2 ORs. Downregulated ORs are chosen less frequently but zonal identity is not affected. 3) Zone-4 and zone-5 neuronal identity has changed and cells physically located in these zones now express markers from other zones. To address these possibilities, we performed Single Cell RNA-Seq (scRNA-seq) comparing swap mice to littermate controls. The chosen OR of each OSN was determined by the RNA counts detected from a given OR gene in each cell (Fig.5A). Zonal identity of the chosen OR was assigned (SFig.5A and SFig.5B). No difference in OR expression levels was detected between the swap and control mice on a single cell level (SFig.5C), suggesting that lower levels of zone-4 or zone-5 OR expression could not be explained by a decrease in expression per cell. In agreement with the skewed OR choice hypothesis, we observe zone-1 and zone-2 population expansion and a dramatic decrease in OSNs choosing ORs from zone-4 and zone-5 (Fig.5A).

**Figure 5.**
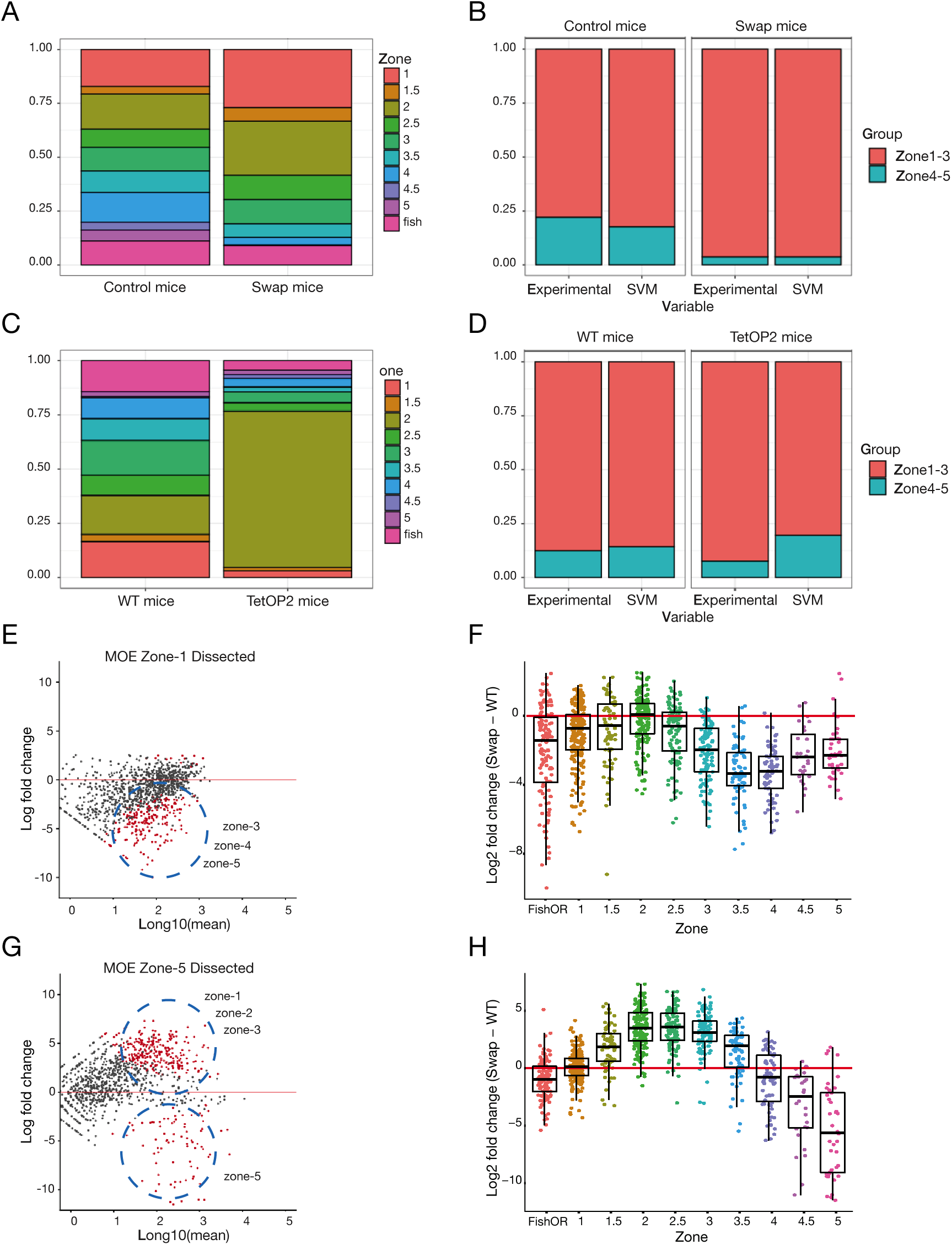
HP1β regulates transcriptional identity and OR choice. A) to D) Single Cell RNA-seq on mature OSNs: A) Stacked bar quantifying fraction of single cells that express an OR from a given zone. Left, littermate control, Right, Swap mice. B) *In silico* prediction analysis (using Support Vector Machine) on transcriptional data from single cell RNA-seq. To train the model, we divided data from control mice in two groups: Cells expressing ORs from zones 1 – 3 and or from zones 4 – 5 (B, first column, experimental). Transcriptional differences (blind to chosen OR information) validate the model (B, second column, SVM). Swap mice analysis shows OSNs express more zone 1-3 OSNs and a change in zonal identity (B, compare 3 and 4 columns). C) Stacked bar quantifying fraction of single cells that express an OR from a given zone. Left, littermate control, Right, gg8tTA;tetO-P2. The tetO forces P2 receptor expression. TetO-P2 allele express a zone-2 OR but maintain their original zonal identity. E) and F) RNA-seq analysis in physically micro dissected cells from zone-1, mOSN sorted cell. Gene expression analysis compering swap mouse versus control mouse. Significantly changed genes are colored red (Padj < 0.05 for greater than 1.5-fold change, Wald test, n = 3). G) and H) RNA-seq analysis in physically micro dissected cells from zone-5, mOSN sorted cells. Significantly changed are colored red (Padj < 0.05 for greater than 1.5-fold change, Wald test, n = 3). Cells from zones-5 show repression from zone-5 receptors and activation of zone-1 to zone-3 receptors.

To determine whether the swap OSNs maintain their transcriptional identity of zone-4 and zone-5 OSNs, even if those OSNs chose a different ORs, we determined the transcriptional differences between OSNs expressing ORs from dorsal zones-1 to zone- 3 and ventral zone-4 to zone-5 (Fig.5B). With the use of Support Vector Machine (SVM) (37) a machine learning tool, we analyzed our scRNA-seq data (Fig.5B). In the control mice, cells defined as mature OSNs were divided into zone-1 to zone-3 and zone-4 to zone-5 expressing cells based on the experimentally chosen ORs, and used to train the model (Fig.5B, control mice, experimental). Roughly 50 mRNAs in addition to chosen ORs differ significantly between the two zonal groups. Some of these transcripts encode proteins involved in axon guidance which is in accordance with the hypothesis that one main function of zonal OR choice is to provide a first layer of organization for axon guidance to the olfactory bulb (38). This analysis reveals that zonal identity at a transcriptional level is associated with the chosen OR. Next, sequencing reads corresponding to the chosen receptors were removed, and the algorithm was used to predict the zonal identity of the given OSN data set (Fig.5B, control mouse, experimental vs SVM), blind to the expressed OR. In the swap mouse, SVM predicts that the ventral zone-4 to zone-5 OSNs have adopted a zone-1 to zone-3 identity (Fig.5B, swap mice, experimental vs SVM), suggesting that these neurons are transcriptionally similar to zone- 1 to zone-3 neurons and their zone-4 to zone-5 transcriptional identity has changed. This result contrasts with a second mouse model (Fig.5C and Fig.5D). An inducible OR, the P2 allele, called tetO-P2iGFP (tetOP2, 39), containing a gg8-tTA driver specific to OSNs. In the gg8tTA;TetOP2 mice, OSNs are artificially biased to express the zone-2 OR *Olfr17* (P2) in the entire epithelium. SVM on a single cell dataset from these mice showed that while the majority of neurons have chosen a zone-2 OR (Fig.5C), the transcriptional zonal identities remain intact (Fig.5D, TetOP2 mice, experimental vs SVM). These data raise the exciting possibility that Hp1β is required not only to ensure zonal OR choice but also to establish zonal transcriptional identity.

To experimentally confirm the SVM prediction, swap and littermate control olfactory epithelia were zonally micro dissected to enrich for zone-1 or zone-5 tissue (Fig.5E and Fig.5F) as previously described (14). Single Cell RNA-seq was performed on pure populations of sorted mature OSNs, using the OMP-ires-GFP allele, from these micro- dissected samples. As previously shown, non-OR genes were nearly unaffected (not shown), while a specific dysregulation of OR genes was observed (Fig.5E). As expected, zone-1 dissected tissue (Fig.5E and Fig.5F) exhibited a drop in OR expression from zone- 3 to zone-5. Surprisingly, in zone-5 dissected tissue (Fig.5G and Fig.5H) the swap mouse recapitulates the drop in zone-5 ORs, but now exhibits an increase in the expression from zone-1 to zone-3 ORs. This increase in zone-1 to zone-3 ORs was masked when using bulk RNA-seq (Fig.4A), likely because ventral zone-5 tissue is a smaller proportion of the epithelium than larger dorsal zones. These results confirm our SVM prediction. OR co-expression was also examined with no clear differences between control and swap mice (SFig.5D). Taken together our single cell RNA-seq data suggest that HP1β regulates OR choice and transcriptional identity establishment that cannot be rescued by HP1α.

### HP1β dependent H3K9me3 methylation participates in OR diversity

HP1β protein and Histone H3 lysine 9 (H3K9) lysine methylation are hallmarks of heterochromatin (21). HP1 has a central part in cell lineage and fate determination (31, 40, 41). H3K9me3 incorporation into the OR genomic clusters occurs as OSNs in specific anatomical zones choose an OR for activation thus defining their identity (14). Given the zonal dysregulation of ORs in the swap mouse, and the gradual incorporation of H3K9me3 into the OR clusters (14), we hypothesized that H3K9me3 levels over OR loci could be zonally altered in the absence of HP1β. To test this, we performed Native Chromatin Immunoprecipitation (nChIP) using an antibody against H3K9me3. Consistent with previous reports (14), we observed differential levels of H3K9me3 incorporation between zone-1 and zone-5 (Fig.6A, control mice). With soft X-ray tomography (SXT), a technique that has the advantage of avoiding PFA to fix interactions, we observe that total heterochromatin levels are higher in tissue from zone-5 vs zone-1, supporting our previous result (Fig.6B). In the absence of HP1β, H3K9me3 levels are reduced over OR loci (Fig.6A, and Fig.6C, swap mice) and total heterochromatin levels are decreased by SXT (Fig.6B). In agreement, our SXT data demonstrate that euchromatin levels increase (Fig.6B). This result shows that an increase in H3K9me3 levels is dependent on HP1β. As a control, we used nChIP to examine the chromatin surrounding the OR enhancers, termed “Greek islands”, as they are regions of euchromatin immersed in heterochromatin. We observe no change in H3K9me3 levels surrounding OR enhancers (SFig.6).

**Figure 6.**
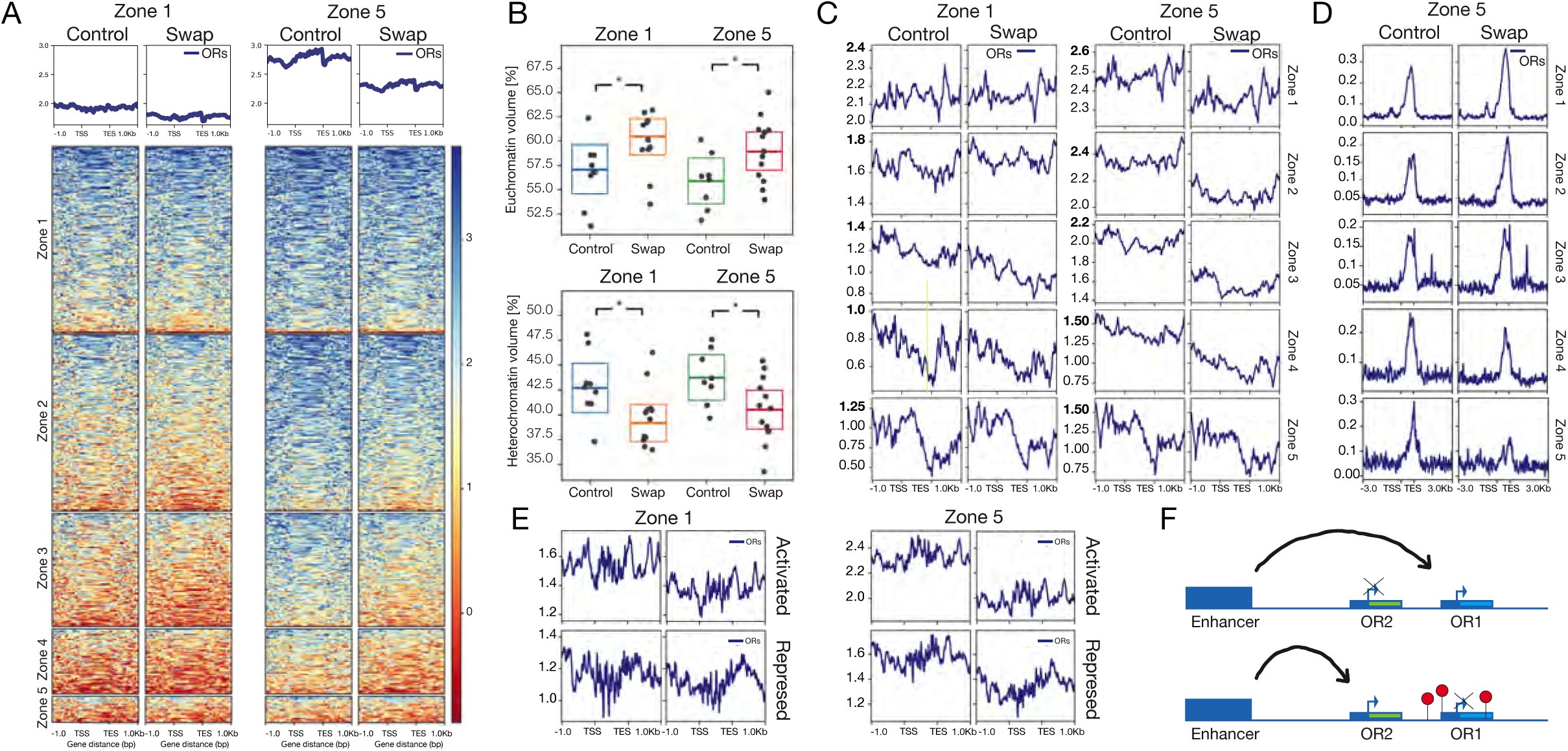
HP1β is implicated in H3K9me3 incorporation and MOE zone formation. A) Native ChIP-seq on mature OSNs from micro dissected tissue from zone-1 and zone-5. Swap versus control mouse shows loss of H3K9 methylation on OR clusters (n = 3 for each genotype). B) Soft X-ray tomography on sorted OMP-GFP + mOSNs from micro dissected zone-1 and zone-5 tissue. Top plot shows percent of euchromatin volume (normalized by total nuclear volu-me) zone1- ctrl (57%) vs zone1-swap (60%) -> *p=0.043, zone5-ctrl (56%) vs zone5-swap (59%) -> *p=0.028. Bottom plot shows percent of heterochromatin volume (normalized by total nuclear volume) zone1-ctrl (43%) vs zone1-swap (40%) -> *p=0.040, zone5-ctrl (44%) vs zone5-swap (40%) -> *p=0.021. C) Native ChIP-seq from micro dissected zone-1 vs zone-5. Reads were bioinformatically separated into zones. Zone-1 to zone-3 exhibit the greatest decrease in H3K9me3. This was observed only on micro dissected cells from zone-5 tissue. D) ATAC-seq on micro dissected zone-1 vs zone-5, ATAC-seq reads were bioinformatically separated into zones (n = 3). E) RNA-seq data sets (figure 4) where bioinformatically segregated into repressed and activa-ted ORs and H3K9me3 levels of incorporation were determined. F) Competition model proposes that OR receptors with a strong promoter will out compete other receptors for their interaction with the enhancer hub. H3K9me3-HP1β dependent methylation will silence strong promoters, giving the opportunity to weaker pro

It is counterintuitive to relate repression with a decrease in heterochromatin. One possible way to explain this discrepancy is with an enhancer competition model (42). In this model (Fig.6F), promoters compete to interact with enhancers. Stronger promoters will interact more frequently for activation. For weaker promoters to be activated, stronger promoters would need to be repressed. We define strong or weak promoters based on their upstream regulatory sequences (see discussion). To gain insight into the feasibility of this model and to explain the partitioning of the OR system into transcriptional zones, we used micro-dissected tissue from zone-1 and zone-5. Next, we bioinformatically segregated OR promoters into the five zones and assessed H3K9me3 enrichment (Fig.6C).

Interestingly, dissected tissue from zone-5 and not from zone-1 shows an H3K9 methylation gradient from zone-5 to zone-1 (Fig.6C, zone-5, control column), showing zone-1 ORs with higher levels of lysine methylation and coincident with zone-5 ORs being active as a consequence of losing methylation. This result supports a model where in tissue from zone-1, low methylation levels favor the expression of stronger promoters. On the other hand, in zone-5 tissue the establishment of gradients of methylation represses strong promoters, enabling “weaker” zone-5 promoters to be chosen.

If HP1β were involved in incorporating H3K9me3 methylation gradients, we would expect to lose those gradients in the swap mouse, which is what we observe (Fig.6C, zone-5 swap mouse). In the absence of HP1β, H3K9me3 levels are reduced over OR loci in all samples from this tissue, but this effect is more pronounced on zone-1 to zone-3 ORs, coincident with a gain in chromatin accessibility as determined by ATAC-seq (Fig.6D). In zone-4 and zone-5 promoters, we observe a mild decrease in H3K9me3 levels (Fig.6C) but a dramatic decrease in chromatin accessibility (Fig.6D), coincident with a loss in activation. These results support a model of enhancer competition where stronger promoters need to be silent to give the chance to weaker promoters to be selected (Fig.6F). With a decrease in heterochromatin levels, even if zone-4 and zone-5 ORs also demonstrate a decrease in H3K9 methylation, their promoters will not be chosen for activation.

As previously shown, distinct gene regulatory programs occur across diverse tissues like the MOE, and the effect in small populations can be masked. For this reason, we decided to examine histone methylation when considering activated vs repressed OR genes (Fig.6E) as determined by RNA-seq (Fig.4A). With this analysis we find that ORs that are activated lose greater levels of methylation compared to repressed ORs (Fig.6E). Remarkably, in the absence of HP1β, promoters from all zones acquire similar levels of H3K9me3 incorporation, losing the methylation gradient observed in control mice that facilitate promoters from all zones to be expressed (Fig.6C, zone-5, control vs swap). Our data support a model where the promoter strengths (68) of dorsal ORs (zone-1 to zone- 3) are able to “out compete” ventral OR promoters. In tissue from zone-5, in a wildtype context strong promoters (zone-1 to zone-3) will be silenced by H3K9me3 to ensure diversity in OR choice.

### Zone-5 OR compartment interactions and nuclear architecture depend on HP1β

The cell nucleus is organized into topologically associated domains that partition the genome into distinct regulatory territories (43). Olfactory sensory neurons form nuclear compartments from the 20 different pairs of chromosomes, making specific and robust interchromosomal contacts among OR gene clusters that increase with cell differentiation as cell identity is established (10, 14, 17). OR compartments from ventral cells (zone-5) are larger and contain a larger amount of H3K9me3 incorporated than dorsal OR genes (zone-1) (14). In *C. elegans* it was recently shown that inactive compartment interactions are dependent on H3K9 methylation levels (44). This complex network of highly specific and stable genomic contacts likely contributes to every step in OSN identity and function, as it regulates both the monogenic and monoallelic olfactory receptor (OR) choice (15) as well as gene expression programs involved in OR processing and OR signaling (45).

Given the properties of HP1 proteins in nuclear architecture organization (32, 33) and the need of H3K9me3 for HP1β to phase separate (46), we decided to examine nuclear architecture of OR gene clusters in the swap mouse. Visual inspection of OSNs in the swap mice revealed fewer strong DAPI dense foci than in controls (Fig.7A). To quantify this, nuclei from swap and control mice were scored as having highly organized chromatin foci or as less organized nuclei, with multiple smaller foci. We found an almost 50% reduction in the number of nuclei with the characteristic “fried egg” shape (Fig.7A). In the control mice 37% of nuclei exhibited this inverted nucleus phenotype, compared to the swap mice with only 20% of nuclei. This result points to a possible defect in OSN nuclear architecture in the swap mouse.

**Figure 7.**
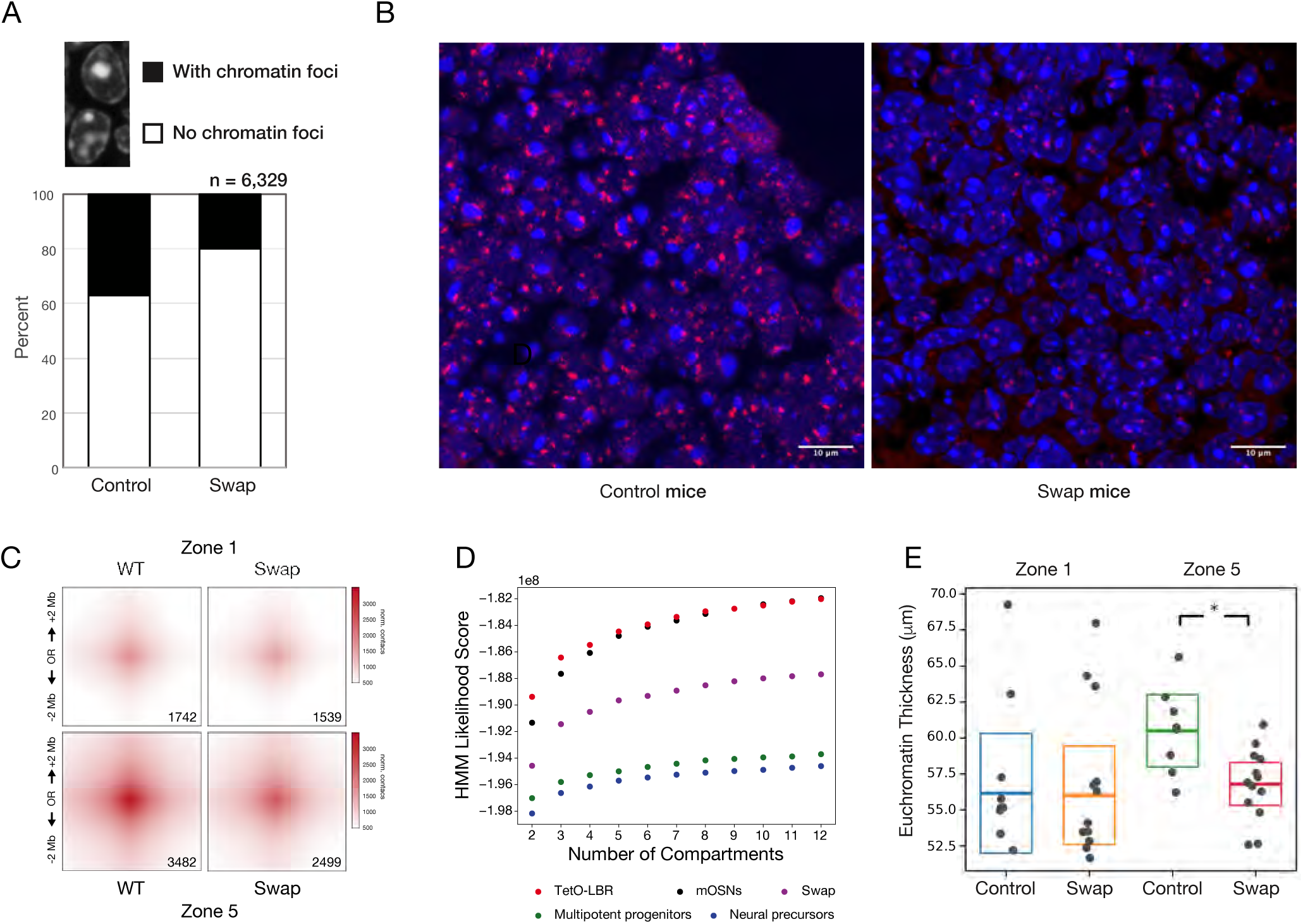
Interchromosomal interactions between OR clusters are affected in the HP1 swap mice. A) Stacked bar plot depicts percent of cells with a chromatin focus, on MOE sections stained with DAPI. B) Representative image from DNA-FISH using a Pan-OR probe. The Pan-OR probe recognizes most of the OR-receptor sequences. Scale bar is 10μm. C) and D) in situ Hi-C experiment on OMP-GFP+ sorted mOSNs from micro dissected zone-1 vs zone-5 (n = 3). C) Aggregated Peak Analysis (APA) matrices of 4Mb regions, shows a decrease in the OR – OR trans contacts, but not in the enhancer – enhancer trans contacts (not-shown). The number located on the right bottom side of each plot represent the number of contacts determined. D) Compartment analysis showed a decrease in the structure of the chromatin compartments compared to mOSNs. Compartment analysis for the Swap mice was compared to mature OSN (TetO-LBR (12) and mOSN) and immature OSN (Multipotent progenitors, Neuronal precursors). E) Euchromatin thickness in control and swap cells. The analysis was performed on sorted mOSN from micro dissected zone-1 and zone-5, imaged using soft X-ray tomography. The analysis showed statistical significance in the euchromatin thick-ness in control and swap cells in zone 5. The statistical significance was evaluated using a two-sided t-test, with *p-value=0.011.

Given that the OR genomic clusters are organized around this DAPI dense focus (12), and are covered in HP1β and H3K9me3 (Fig.1, SFig.2, Fig.6A), we hypothesized that the OR loci would be disorganized in the swap mouse. To test this, we performed DNA-FISH using the Pan-OR probe (Fig.7B), which recognizes many olfactory receptor gene loci. Control Pan-OR signal was readily observed surrounding the heterochromatic focus in control mice, whereas in the swap mouse, Pan-OR signal was reduced, fewer puncta were observed per cell, and they were not organized around the heterochromatic focus/foci. The analysis of 3-dimensional data of the nuclei collected using SXT (47) also shows that chromatin foci break into smaller heterochromatin dense domains (SFig.7A), revealing a loss of OSN characteristic nuclear architecture. Bashkirova et al (14) describe characteristic and distinct compartment interactions when comparing zone-1 versus zone-5. A change in identity as we report should also mean a change in nuclear architecture. To answer this question, we performed *in situ* Hi-C (16) on pure sorted mature OSN populations from micro dissected zone-1 and zone-5 tissue from the swap and control littermates (Fig.7C). Aggregated peak analysis (APA) confirms as previously reported that zone-5 ORs exhibit more interchromosomal interactions than zone-1 ORs. Interestingly, in the dissected zone-5 swap mice, interchromosomal interactions go down by 28%, homogenizing the differences that distinguish zone-1 ORs from zone-5 ORs (Fig.7C). SXT confirmed this result (SFig.7A and SFig.7B), as the relative composition of pericentromeric chromatin shows a reduction in size. Compartment analysis on Hi-C data (Fig.7D) confirm less structured chromatin clusters in the swap mouse versus control littermates. These results suggest that in the absence of HP1β, zone-5 OSNs exhibit homogenized H3K9me3 levels and chromatin architecture, breaking the epigenetic footprint that differentiates ORs across zones.

As cells differentiate within an organism, genes that are no longer expressed will become silenced (28), forming blocks of heterochromatin that separate active genes from repressed genes (19). HP1 proteins have been suggested to partition constitutive heterochromatin (32, 33, 49). We hypothesized that zone-5 nuclear architecture should have a better separation of heterochromatin vs euchromatin. If this hypothesis were true, we should observe a less organized distribution of chromatin in the nuclei of the swap mouse. To test this hypothesis, we used SXT to determine the euchromatin thickness within the nucleus (Fig.7E). This technique can tell us if heterochromatin versus euchromatin separation is less homogenous in a zonal manner. As expected, WT OSNs from zone-5 OR show a clearer separation of euchromatin from heterochromatin. In the swap mouse the differences found between zone-5 and zone-1 tissue are erased. These results support a model where HP1β is implicated in nuclear chromatin organization, partitioning euchromatin from heterochromatic regions, facilitating the recruitment of ORs from more dorsal zones into heterochromatic compartments that will ensure their silencing in ventral zones.

### HP1α fails to form OR clusters around the heterochromatin foci

If HP1α and HP1β are highly conserved paralogs, why is HP1α unable to rescue the HP1β KO phenotype? We envision three different scenarios to respond to this question. First, HP1α does not interact with the OR clusters. Second, HP1α fails to segregate OR clusters from pericentromeric chromatin and third, HP1α fails to recruit the enzymatic activities necessary for zonal OR choice. RNA-seq shows HP1α is not present in mOSN cells (Fig.1A). To understand if this was simply a problem of availability, we forced the expression of HP1α in OSNs (Fig.8A). With the use of the swap mouse that includes the tagged HP1α knock-in coding sequence under the endogenous HP1β promoter and the use of an antibody against the tags, we found HP1α is enriched at the chromatin foci at early stages of differentiation, but in mature OSNs, HP1α stains diffusely through the nucleus and heterochromatic foci. This result suggests that HP1α fails to be enriched in the heterochromatin clusters surrounding the heterochromatin foci. In contrast by ChIP-seq, we observe HP1α interacting with the OR clusters in mOSNs (SFig.8). This suggests that in the absence of HP1β, HP1α is capable of interacting with the OR clusters. Previous reports have shown that *in vitro,* the addition of HP1β will dissolve HP1α-phase transition droplets (50). Visual inspection shows absence of HP1α enrichment at the periphery of the chromatin foci. To confirm this, we used SoRA high-resolution microscopy to quantified the radial profiles of WT HP1β, WT HP1α, and swap HP1α protein localization within the DAPI dense foci. We found that swap HP1α does not migrate to the edges of OR chromatin foci (Fig.8B) as HP1β does. This result, in addition to the Hi-C data (Fig.7C and Fig.7D) and SXT (Fig.7E and SFig.7A) shows that HP1α cannot organize the OR clusters into discrete OR compartments compared to the control mice.

**Figure 8.**
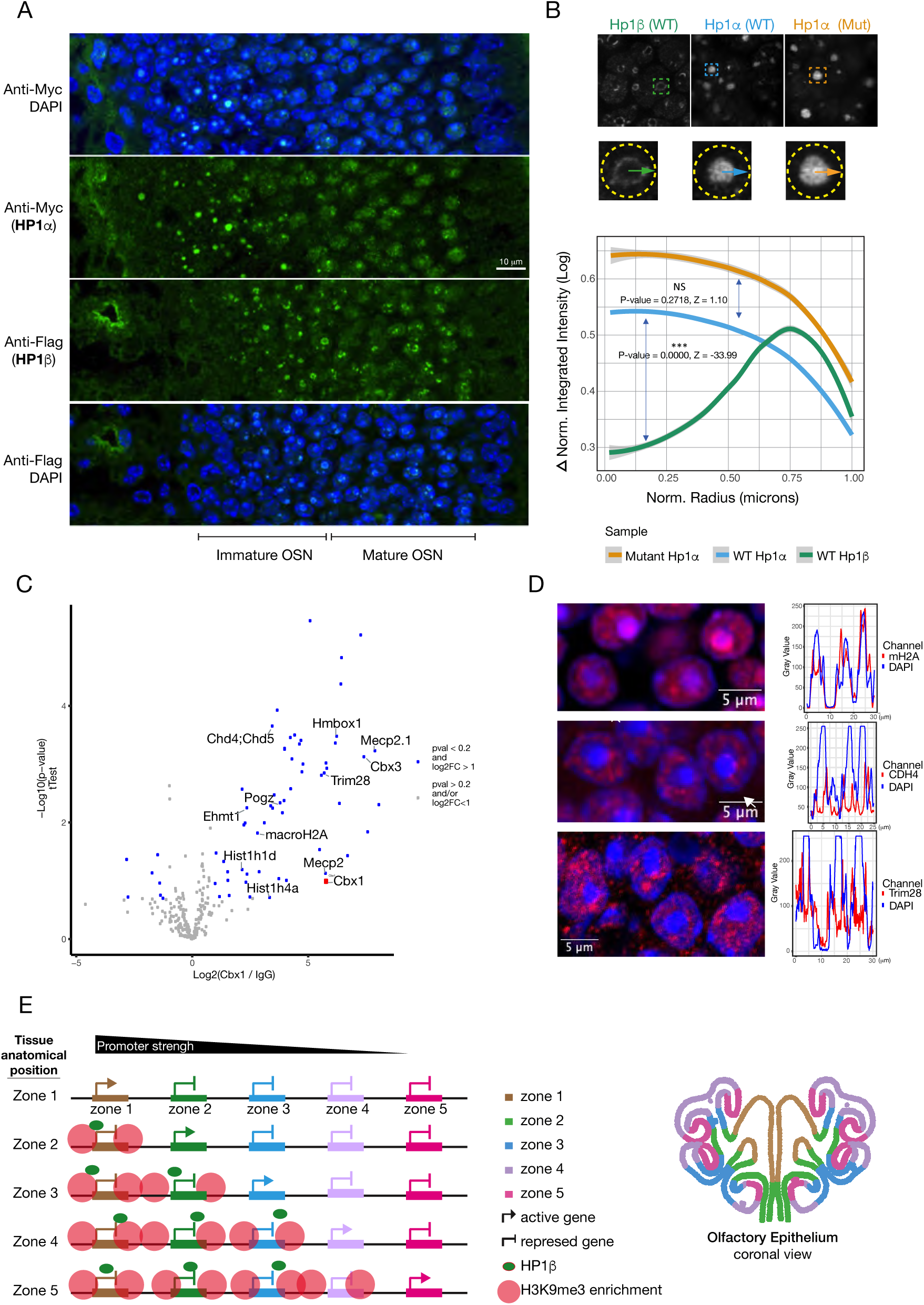
HP1β and not HP1α is recruited to the OR clusters with chromatin remodeling activities. A) IF against HP1β-Myc tag or HP1α-Flag tag shows HP1α fails to localize with heterochromatin in mOSNs. Scale bar is 10μm. B) Radial profile analysis for the proteins Hp1α and Hp1β in DAPI dense foci of the MOE. A circle was drawn around each focus, and the radial profile was measured. See material and methods for details. C) Mass Spectrometry on proteins immunoprecipitated from MOE tissue using an specific antibody against HP1β. D) IF imaging on MS candidates and staining intensity determination using FIJI and Intensity geomline. Scale bar is 5μm. E) Working model. H3K9 methylation is incorporated at different levels in the mouse MOE. H3K9me3 shows gradients of repression that decorate the 5 anatomical zones. We suggest methylation will silence stronger promoter compared to the receptors expressed in the corresponding zone, giving a chance to weaker receptors to be chosen. The recruitment of HP1β will come with chromatin remo-deling activities and repression. Antibodies used on IF: mH2A (ab208879), CDH4 (ab70469) and Trim28 (ab22553).

We also hypothesized that the protein interacting partners of these two Hp1 proteins could explain HP1α’s inability to incorporate into or recruit H3K9me3 to OR clusters. With the use of Mass Spectrometry (MS) we determined the proteins interactors for HP1α and HP1β (Fig.8C and SFig.8B). We performed three replicates for each immunopurification, with antibodies against HP1α, HP1β and a control IgG using a mix of Cbx and WT MOE lysate. Pearson correlation analysis shows high correlation between replicates and distinct separation between IP targets (not shown). Gene set enrichment analysis (GSEA) shows enrichment of proteins involved in histone binding, histone deacetylase binding, among others, for both proteins (SFig.8C). Amongst the top candidates for HP1β (cut off pval < 0.05 and log2FC > 2), we identify previously reported interactors, including Trim28 (51), Ehmt1 (52, 53), macroH2A (54), and Chd4 (55). Some of these candidates are of particular interest including, Ehmt1 a histone methyltransferase, involved in H3K9me2 incorporation and a HP1β interactor (52, 53). The Trim family of proteins has previously been shown to recruit HP1 proteins to diverse genomic regions (56, 57). With the use of IF and dependent on the availability and quality of antibodies, we confirmed the colocalization of selected MS candidates in HP1β compartments (Fig8.D, SFig.8D). Interestingly, macroH2A was identified as an HP1β interactor, which fits with the observation that its incorporation into chromatin happens in parallel with HP1β recruitment as OSNs differentiate (Fig.8D and SFig.8D). Remarkably, macroH2A staining colocalizes with the DAPI foci staining (Fig.8D), but it also gets enriched in a halo around the chromatin foci, as we observe for HP1β in mOSNs (Fig.6B). The Chromodomain-Helicase-DNA-Binding Protein 4 (CDH4) stains discrete domains around the OR foci (Fig.8E). CDH4 is an ATP-dependent helicase that binds and distorts nucleosomal DNA (58, 60) and acts as a component of the histone deacetylase NuRD complex which is involved in chromatin remodeling and repression (58, 61, 62, 63). CDH4 enrichment in chromatin foci and its deacetylase activities coincide with the gain of repressive chromatin on the OR foci as described by Clowney et al. (12). Finally, the nuclear corepressor for KRAB domain-containing zinc finger proteins (KRAB-ZFPs, also known as Trim28) is known for mediating transcriptional repression by recruiting repressive complexes (64, 65). Trim28 staining in the MOE is diffuse through the epithelium (SFig.8D) but stains around the nuclear foci in mature OSNs (Fig.8D). In conclusion, Hp1β recruitment to the chromatin foci is accompanied by chromatin remodeling proteins that could facilitate the incorporation of repressive chromatin. Taken together, these data lead us to propose that HP1β but not HP1α, through its interaction with the OR foci and specific protein interactors, recruits chromatin remodeling complexes, and facilitates H3K9me3 incorporation into the OR clusters.

## Discussion

In this work we propose a model for OR receptor gene choice and establishment of zonal identity that depends on the competition of OR promoters to interact with the OR enhancer hub. We propose that this competition is mediated by the incorporation of H3K9me3 in an HP1β dependent manner (Fig.8E). Our model proposes that the incorporation of H3K9me3 will create repression gradients, blocking strong promoters from expression in all the zones. The competition of promoters for enhancers would depend on the promoter strength defined by their sequence composition. In dorsal zones from the MOE, where low levels of H3K9me3 were found (Fig.7), we propose that strong promoters (zone-1 receptor) will interact with the OR enhancer hub and become active. In ventral zones from the MOE, were receptors from zone-5 have the propensity to be chosen, receptors from zone-1 to zone-4 harbor higher levels of histone methylation (Fig.7), giving a chance to weaker promoters (zone-5) to be chosen.

Monoallelic regulation of OR expression has been explained by two main models. The stochastic model suggests that the same set of transcription factors (TF) will participate in the stochastic selection of one allele (39). The deterministic model proposes the use of specific combinations of *cis*-elements and transcription factors to express a gene within a defined spatial zone (81). As the deterministic model proposes, diverse OR gene promoter structures have been described, with receptors regulated by TATA-box (29), or TATA-less containing promoters (79). Transcriptome sequencing shows differences in the strength of these TATA-boxes (68). Some promoters were reported to contain an EBF-like site, but not others. Finally, in a recent publication comparing the variability of gene expression between dorsal versus ventral receptors (DV-score), a specific score for each OR receptor was determined that will define the spatial location at which the specific receptor will be expressed (80). The DV-score depends on the genomic sequences located upstream of each receptor and on the level of heterochromatinization (80). We support the coexistence of the two models: a deterministic mechanism founded on a range of regulatory sequences that will generate a repertoire of promoters with different strengths (similar to the DV-score) and a stochastic mechanism that will stochastically choose among promoters having the same strength. Histone H3K9 methylation will silence strong promoters, thus allowing weaker promoters to be chosen. With this work we reveal a function for HP1β in OR gene choice and neuronal diversity. Our swap mouse design allowed us to test the function of HP1 proteins *in vivo*, enabling imaging and biochemical assays in the context of neuronal differentiation and cellular identity acquisition. Importantly, we uncovered a mechanism by which a stochastic process becomes biased.

### HP1β has a specific role in OR receptor choice, that HP1α cannot rescue

HP1β protein was found to colocalize with the heterochromatic chromatin foci (12) and as an H3K9me3 interactor (66), with a role in diverse processes including nucleating repressive chromatin (19) and gene silencing (67). With the use of IF and RNA-Seq (Fig.1), we showed the dynamics of HP1α and HP1β localization and function through differentiation, OR choice, and neuronal maturation. These proteins are expressed during different windows of neuronal differentiation, suggesting distinct roles in the process of OR selection. Furthermore, HP1α expression decreases early during differentiation, suggesting either an early function for this protein or a specific role for HP1β. In addition, HP1α and HP1β were found to occupy different chromatin compartments in OSNs and progenitors, reflecting separate functions in the process of OR choice.

By using an HP1β constitutive KO model, we found HP1β participates in the process of OR choice (Fig.2). Our swap model system confirms these findings by RNA-Seq (Fig.4A) and using a specific antibody against an OR from zone-5 (Fig.4C). We show Cbx5 knock-in (HP1α) expression rescues the high lethality observed in the HP1β KO, however mutant mice are smaller and rarely survive past three weeks after birth. Taken together, we show for the first time *in vivo* that HP1β has specific functions on transcriptional regulation that HP1α cannot rescue.

### HP1β regulates OR gene choice and transcriptional identity

The HP1β KO mice and the swap mice showed skewed OR expression to receptors from zones-1 to zone-3. ORs from ventral zones were less likely to be chosen in both genetic contexts (Fig.2D, Fig.4D). Single Cell RNA-Seq confirms that fewer OSNs choose zone-4 or zone-5 ORs (Fig.5A). Comparative expression analysis shows no drop on transcriptional OR levels, supporting an effect related to OR choice instead of OR transcriptional levels. *In silico* analysis from single cell RNA-Seq data (Fig.5B) and RNA-Seq on micro dissected tissue from zone-1 and zone-5 OSNs (Fig.5E to Fig.5H) showed that cells have chosen a receptor from a different zone. Transcriptional identity of zone-5 OSNs is skewed to zone-1 to zone-3 receptor identities. These results demonstrate that the absence of HP1β affects not only OR choice, but also cell identity, as receptors from dorsal zones of the main olfactory epithelium have colonized the tissue.

### HP1β regulates OR choice by a mechanism that includes H3K9me3

The HP1β protein is known for its participation in chromatin formation and gene silencing, through mechanisms that involve chromatin methylation (7). In the swap mouse H3K9me3 levels go down globally including over OR gene clusters (Fig.6). Curiously, the effect on H3K9me3 is not homogenous, dorsal ORs that incorporate heterochromatin first (described in 14) exhibit a much more severe reduction in H3K9me3. This result supports H3K9me3-HP1β dependent incorporation in ORs via a regulated mechanism to be described. In tissue from zone-5 but not from zone-1, we found that H3K9me3 forms gradients of repression which serve to differentiate ORs from different zones (Fig.6C). We suggest that the recruitment of histone methyltransferases and chromatin remodeling complexes by HP1β including G9a and the NuRD complex identified in this work as an HP1β interactor (Fig.8C), facilitates the incorporation of H3K9me3, blocking OR receptors with stronger promoters. H3K9me3 will play a role in restricting the repertoire of receptors from which each neuron can choose, parsing the 1,300 OR receptors into zonal subsets.

### HP1β Nuclear Architecture

The process of OR gene choice requires the complete reorganization of the OSN nuclear architecture, which occurs during neuronal differentiation (12, 47). During this process, OR genes from different chromosomes converge into specialized nuclear bodies and chromatin compartments, that may mediate both their efficient silencing and their robust transcriptional activation (12, 14). The aggregation of OR loci is synchronous with the incorporation of repressive chromatin. ORs from zone-4 and zone-5 were shown to be the last group of OR genes to be incorporated into the repressive chromatin foci (14), and to harbor lower levels of heterochromatin when compared to ORs from other zones (14).

We propose a role for HP1β in incorporating H3K9me3 and facilitating OR gene recruitment into the nuclear chromatin foci. This could be a direct role of HP1β or as a consequence of changes in histone methylation levels. From our work and work from other labs, we suggest HP1β and HP1α have roles in organizing the chromatin foci. *In vitro* studies have shown that HP1β can form liquid droplets in the presence of H3K9me3 (48), which may explain the formation of the OR compartments around the nuclear foci decorated with HP1β and H3K9me3. We showed that HP1α and HP1β segregate into different chromatin compartments, even at the interior of an immature heterochromatic focus (Fig1.E, SFig.1B). HP1α exhibits a dramatic re localization from widespread nuclear staining in basal cells to pericentromeric chromatin foci in iOSNs (Fig1.E, SFig.1C). This coincides with the recruitment of HP1β and H3K9me3 to the nuclear foci. Our imaging analysis suggests that HP1β, likely with OR gene clusters, migrates from the interior of the nuclear focus to form a ring at the exterior of the focus. This shift in localization could be explained by the incorporation of H3K9me3 and the properties of phase transition. These structures are partially lost in the absence of HP1β (Fig.7A to Fig.7C, Fig.8B). We suggest that the relationship between HP1β and HP1α may regulate OR segregation into chromatin foci and chromatin organization. In the absence of HP1β the euchromatic thickness is reduced, illustrative of less organized nuclei (Fig.7E). Compartment analysis of Hi-C data confirms less structured chromatin clusters (Fig.7D) and 30% of the interchromosomal interactions are lost (Fig.7C). Radial profile analysis of imaging data shows HP1α does not accumulate around the heterochromatic foci as HP1β does, affecting the ability of OR clusters to migrate to the chromatin foci periphery (Fig.8B). While additional work is required to confirm this model, our data suggest that HP1 proteins play an important role in organizing nuclear architecture in the MOE.

In this work we propose a role for HP1β in incorporating gradients of H3K9me3, partitioning the MOE in zones, with differences in their chromatin architecture that facilitate OR choice of the complete repertoire of ORs. We demonstrate a specific role for HP1β and not HP1α in OR choice and transcriptional identity formation.

## Acknowledgements

We thank Dra Mayra Furlan-Magaril and Dr Nora Elphege-Pierre for helpful feedback on the manuscript. We acknowledge Ira Schieren and the Flow Cytometry Core at Columbia University. MJ would like to acknowledge funding from NIH/NIGMS (R35GM152258).

## Author Contributions

Martín Escamilla del Arenal, conceived and designed the study, performed experiments, participated in swap mouse design and wrote manuscript. Rachel Duffié designed, performed, and analyzed single cell RNA-Seq, IF, HR microscopy, as well participated in manuscript writing. Hani Shayya, analyzed single cell data. Valentina Loconte, Axel Ekman and Carolyn Larabell performed SXT and analyzed SXT data. Lena Street and Marko Jovanovic performed MS. Valentina Loconte and Lena Street provided helpful feedback on manuscript. Adan Horta performed Hi-C compartment analysis. Daniele Canzio facilitated SXT study. Kevin Monahan participated in manuscript writing. Stavros Lomvardas designed swap mouse and provided financial support.

## Material and Methods

### Mouse Strains

Mice were treated in compliance with the rules and regulations of IACUC under protocol number AC-AAAT2450 and AC-AABG6553. Mice were sacrificed using CO2 following cervical dislocation. Cbx KO mice are from EUCOMM, Sanger (strain name: EPD0027_2_B01). Gene: Cbx1, colony prefix: MAAT, ESC clone ID: EPD0027_2_B01, allele: *Cbx1^tm1a(EUCOMM)Wtsi^*, allele type: Knockout First, Reporter-tagged insertion with conditional potential. Cbx Swap mouse was custom designed and made by Biocitytogen (Regeneron Pharmaceuticals), Gene: Cbx1, allele: Cbx1(Cbx5) CKI, model design: Cbx1^tm1a(EUCOMM)Wtsi^. The targeting vector containing the construct LoxP, flag-Cbx1, STOP, myc-Cbx5, LoxP was inserted in Cbx1 locus by homologous recombination, eliminating exons 2 to 5 from the endogenous Cbx1 coding sequence (gene ID: 12412 (NCBI)).

### Fluorescence-activated cell sorting

Cells from the MOE were dissociated with papain for 40 minutes at 37°C according to the Worthington Papain dissociation protocol. Cells were washed 2X with cold PBS, passed through a 40-μM strainer filter. For RNA-seq and native ChIP-seq, DAPI-negative (live cells), fluorescent cells (either +Ngn-GFP or +OMP-GFP) were collected. For Hi-C, cells were fixed 10 minutes in 1% formaldehyde in PBS at RT. The reaction was quenched by adding glycine, and cold PBS washed before sorting fluorescent cells.

### DNA FISH

Fresh mouse olfactory epithelium was embedded in Optimal Cutting Temperature compound (OCT - Sakura) and frozen to -20C. Cryosections were cut at 6μM, air-dried for at least 30 minutes and fixed in ice-cold 4% PFA for 5 min. Fixation was stopped by rinsing the sections 2 times with permeabilization media PBS 1% Triton (PBST). DNA was fragmented with 0.1M Hydrochloric Acid (HCl) for 15 minutes at Room Temp (RT), digested with 3 mg/ml of RNase A (Thermo Scientific™, Catalogue No. 10753721) in PBST for 1 hr at 37°C and dehydrated by 3X 70%/90%/100% ethanol washes at 45C for 5 minutes. Sections were denatured at 85°C for 5 min in 2xSSC, 75% formamide (Invitrogen), and immediately dehydrated as before but with ethanol at ice-cold temperature. PanOR probe (Biotin-Nick Translation Mix, Roche, Cat. Number 11745824910) was added at 25 ng/μL and covered with 8mm circular coverslips, sealed with rubber cement and incubated O/N 37C. Slides were washed in 2xSSC, 55% formamide, 0.1% NP-40 3X for 15 minutes, rinsed in PBST, blocked in TNB (Promega TSA kit), and incubated 2 hr RT with anti-dig conjugated to DyLight fluors (Jackson Immunoresearch), 3X washed in PBST + 8% formamide and mounted. Images were collected on a confocal Zeiss LSM700 microscope.

### Immunofluorescence and Microscopy

Olfactory epithelium was dissected and frozen in OCT compound as described above. Cryosections were collected at 12μM with control and experimental samples on the same slide to reduce variability across genotypes. Slides were air dried for 10 minutes, fixed in a slide mailer with RT 4% PFA; 1X PBS for 5 minutes, and washed 3X with 1X PBS. Slides were blocked in a slide mailer with blocking solution (1X PBS, 2% Donkey Serum (Sigma), and 1% Triton X-100) for 30 minutes. Primary antibodies were diluted 1:200 in blocking solution and slides were incubated overnight in a humid chamber with primary antibody solution, covered with a coverslip to prevent evaporation. The following day, coverslips were displaced with excess PBS 1X, and slides were washed with PBS 1X; 0.1% Triton X-100 3x5 minutes in a slide mailer. Secondary antibodies (Jackson Immunoresearch and Invitrogen) were diluted 1:500 and 1mg/mL DAPI was diluted 1:1000 in blocking solution. Secondary antibody solution was applied to slides and covered with a coverslip and slides were incubated in a humid chamber for 45 minutes at RT. Coverslips were washed off with excess PBS 1X, and slides were washed 3x5 minutes with 1X PBS; 0.1% Triton X-100. Slides were mounted with Vectashield antifade mounting medium (Vector Laboratories) and sealed with nail polish. Images were taken on a Zeiss LSM 700 Confocal microscope using a 40 or 63X objective, or with a SoRa- W1-Yokogawa spinning disk confocal microscope. High resolution images were collected with a z-step size of 0.1 μM, and stacks were deconvolved using the FIJI plug-in Microvolution. Image quantification was performed in FIJI and R.

### Radial profile analysis of high-resolution microscopy DAPI dense foci

Imaging data were collected from 197 HP1α -positive foci in WT mice, 223 HP1β -positive foci in WT mice, and 302 HP1α -positive foci in swap mice. For each fluorescent focus, the radial distance from the center was normalized to 1. Normalized Integrated Intensity (measure of HP1α or HP1β fluorescence intensity) values were log-transformed. For each focus, the change in log-transformed normalized intensity (Δ) was calculated using the brightest and dimmest value for each focus. This analysis examines the relative change in intensity within each focus, rather than on absolute fluorescence intensity differences among the three experimental groups.

Radial profiles were plotted using a locally estimated scatterplot smoothing (loess) regression, with 95% confidence intervals displayed for each group. Statistical comparisons of Δ (log-transformed) normalized intensity as a function of normalized radius were performed using Fisher’s Z transformation (Fisher, 1925) to assess whether correlation profiles differed significantly between groups: HP1α WT vs. HP1β WT, and HP1α WT vs. HP1α swap.

### RNA-Seq

Fresh mouse olfactory epithelia were used for RNA-Trizol extraction using standard protocols in an RNA safe area. DNase treatment was performed with Turbo DNA-free kit (Ambion, Cat. Number AM1907), QC and sequencing libraries were prepared with Nugen NuQuant RNA-seq library system and sequenced on a HiSeq2500. Cut adapt was used to remove adapter sequences and reads were aligned to the mm10 genome with STAR. Samtools was used to select high mapping quality reads (-q 30). Normalization, calculation of FPKM (converted to TMP), and differential expression analysis was performed in R with DEseq2. For all RNA-seq data p-values refer to adjusted p-value (padj) calculated in DEseq2. RNA-Seq data from sorted cells during neuronal differentiation have been described in (82) and are posted in the GEO repository GEO: GSE271029.

### Native Chromatin Immunoprecipitation

Native ChIP was performed as described in Magklara et al. (2011). Briefly, nuclei isolation: FACS-sorted cells were pelleted (600 x g, 10 min, 4°C) and resuspended in ice-cold Buffer I (0.3M Sucrose, 60 mM KCl, 15 mM NaCl, 5 mM MgCl2, 0.1 mM EGTA, 15 mM Tris-HCl pH 7.5, 0.1 mM PMSF, 0.5 mM DTT, 1x protease inhibitors). Cells were lysed by adding an equal volume of ice-cold Buffer II (Buffer I + 0.4% NP-40) and incubating on ice for 10 minutes. Nuclei were pelleted (1000 x g, 10 min, 4°C) and resuspended in 250 µL of ice-cold MNase buffer (0.32M Sucrose, 4 mM MgCl2, 1 mM CaCl2, 50 mM Tris-HCl pH 7.5, 0.1 mM PMSF, 1x protease inhibitors). Micrococcal Nuclease (MNase) Digestion: 0.1 U of MNase (Sigma) per 100 µL of MNase buffer was added, and the mixture was incubated for 1 minute and 40 seconds in a 37°C water bath. Digestion was stopped by adding EDTA to a final concentration of 20 mM. Chromatin Fractionation: The first soluble chromatin fraction (S1) was pelleted and supernatant was stored overnight at 4°C. The undigested material (pelleted) was resuspended in 250 µL of ice-cold Dialysis Buffer (1 mM Tris-HCl pH 7.5, 0.2 mM EDTA, 0.1 mM PMSF, 1x protease inhibitors) and rotated overnight at 4°C. The undigested material was pelleted by centrifugation and constitute S2 fraction. S1 and S2 fractions were combined and 5% of the combined chromatin was saved as input. Immunoprecipitation (IP): Chromatin was diluted to 1 mL in Wash Buffer 1 (50 mM Tris-HCl pH 7.5, 10 mM EDTA, 125 mM NaCl, 0.1% Tween-20, 5x protease inhibitors) and rotated overnight at 4°C with 1 µg of antibody. Dynabeads (10 µL Protein A and 10 µL Protein G per IP) were pre-blocked overnight at 4°C with 2 mg/mL yeast tRNA and 2 mg/mL BSA in Wash Buffer 1. Blocked beads were washed once with Wash Buffer 1, then added to the antibody-bound chromatin and rotated for 2-3 hours at 4°C. Beads were washed 4x with Wash Buffer 1, 3x with Wash Buffer 2 (50 mM Tris-HCl pH 7.5, 10 mM EDTA, 175 mM NaCl, 0.1% NP-40, 1x protease inhibitors), and 1x with TE buffer (pH 7.5). Elution and DNA Purification: Immunoprecipitated DNA was eluted by resuspending the beads in 100 µL of Native ChIP Elution Buffer (10 mM Tris-HCl pH 7.5, 1 mM EDTA, 1% SDS, 0.1 M NaHCO3) using a thermomixer (37°C, 900 rpm, 15 min). This elution step was repeated twice, and the eluates were combined. Library Preparation and Sequencing: Sequencing libraries were prepared using the NuGEN Ovation V2 DNA-Seq Library Preparation Kit. Sequencing was performed using 50 bp paired-end (PE) reads on a HiSeq 2500 or 75 bp PE reads on a NextSeq 550.

### ATAC-seq

ATAC-seq was performed from live sorted cells using the protocol developed by Buenrostro et al., 2015. Briefly, cells were pelleted and then resuspended in lysis buffer (10 mM Tris-HCl, pH 7.4, 10 mM NaCl, 3 mM MgCl2, 0.1% IGEPAL CA-630). Nuclei were immediately pelleted (1000 rcf, 10 min, 4°C). Pelleted nuclei were resuspended in transposition reaction mix prepared from Illumina Nextera reagents (for 50 μL: 22.5 μL water, 25 μL 2xTD buffer, 2.5 μL Tn5 Transposase). The volume of the Tn5 transposition reaction was scaled to the number of cells collected: 1 μL mix per 1000 cells. If fewer than 10,000 cells were collected by FACS, 10 μL scale reactions were performed. Transposed DNA was column purified using a Qiagen MinElute PCR cleanup kit (Qiagen). The transposed DNA was then amplified using barcoded primers and NEBNext High Fidelity 2x PCR MasterMix (NEB). Amplified libraries were purified using Ampure XP beads (Beckman Coulter) at a ratio of 1.6 μL of beads per 1 μL of library and eluted in 30 μL of elution buffer (10 mM Tris-HCl pH 8, 0.1 mM EDTA).

### ChIP-seq and ATAC-seq Analysis

Adapter sequences were removed with CutAdapt. ChIP-seq and ATAC-seq reads were aligned to the mouse genome (mm10) using Bowtie2 (Langmead and Salzberg, 2012). Default settings were used, except a maximum insert size of 1000 (-X 1000) was allowed for ATAC-seq and native ChIP-seq data since these data sets contained some large fragments. PCR duplicate reads were identified with Picard and removed with Samtools (Li et al., 2009). Samtools was used to select uniquely aligning reads by removing reads with alignment quality alignments below 30 (-q 30). HOMER (Heinz et al., 2010) was used to call peaks of ChIP-seq signal using the ‘factor’ mode and an input control. Consensus peak sets were generated by selecting peaks that overlapped between biological replicates and extending them to their combined size. For signal tracks, replicate experiments were merged, and HOMER was used to generate 1 bp resolution signal tracks normalized to a library size of 10,000,000 reads. For H3K9me3 ChIP-seq replicate experiments were merged, and HOMER was used to generate 1 bp resolution signal tracks normalized to a library size of 10,000,000 reads. Regions enriched for H3K9me3 were identified by running HOMER peak calling.

### Zonal dissection of the olfactory epithelium

Fluorescent signal in Olfr545-delete-YFP (zone 1 OR), Olfr17-ires-GFP (zone 2 OR), and Olfr1507-ires-GFP(18) (zone 5 OR) mice were used to standardize dorsal(zones-1) MOE and ventral (zone 4-5) MOE micro dissections (See Bashkirova et al. 2023). Accuracy of dissections was confirmed by RNA-seq.

### Single cell RNA-seq in olfactory lineage cell types

Fresh mouse olfactory epithelium was dissected and dissociated with papain for 40 minutes at 37°C according to the Worthington Papain Dissociation System. Cells were washed 2x with cold PBS before passing through a 40-μM strainer. Library preparation and Sequencing was conducted by the Columbia University Genome Center. Experiment was performed in biological replicate. Data were aligned using Cell Ranger (10X Genomics) and analyzed using Seurat.

### In situ Hi-C

In situ Hi-C and library preparation was performed as described (14). Briefly, FAC-sorted cells (inputs ranged from 150,000 to 500,000 cells) were pelleted at 500 rcf for 10 minutes and lysed in Lysis buffer (50 mM Tris pH 7.5 0.5% NP40, 0.25% sodium deoxychloate0.1% SDS, 150 mM NaCl and 1x protease inhibitors) by rotating for 20 min at 4°C. Nuclei were pelleted at 2500 rcf, permeabilized in 0.05% SDS for 20 min at 62 °C, then quenched in 1.1% Triton-X100 for 10 min at 37 °C. Nuclei were then digested with DpnII (6U/μL) in 1×DpnII buffer overnight at 37 °C. In the morning, nuclei were pelleted at 2,500g for 5 min and buffers and fresh DpnII enzyme were replenished to their original concentration and nuclei were digested for 2 additional hours. Restriction enzyme was inactivated by incubating 20 minutes at 62 °C. Digested ends were filled in for 1.5 hours at 37 °C using biotinylated dGTP. Ligation was performed for 4h at room temperature with rotation. Nuclei were pelleted and sonicated in 10 mM Tris pH 7.5, 1 mM EDTA, 0.25% SDS on a Covaris S220 (16 minutes, 2% duty cycle, 105 intensity, Power 1.8-1.85 W, 200 cycles per burst, max temperature 6°C). DNA was reverse crosslinked with RNAse A and Proteinase K overnight at 65 °C then purified with 2× Ampure beads following the standard protocol and eluted in water. Biotinylated fragments were enriched with Dynabeads MyOne Strepavidin T1 beads and on bead library preparation was carried out with NuGEN Ovation V2 DNA-Seq Library Preparation Kit, with some modifications: instead of heat inactivation following end repair beads were washed 2x for 2 min at 55 °C with Tween Washing Buffer (TWB)(0.05% Tween, 1 M NaCl in TE pH 7.5) and 2x with 10 mM Tris pH 7.5 to remove excess detergent. After ligation of adapters, beads were washed 5x with TWB and 2x with 10 mM Tris pH 7.5. Libraries were amplified for 10 cycles and cleaned up with 0.8V Ampure beads. Each experiment was performed with two biological replicates and prepared Hi-C libraries were sequenced 75PE on NextSeq 500.

### In situ Hi-C analysis

Reads were aligned to the mm10 genome using the distiller pipeline (https://github.com/mirnylab/distiller-nf), uniquely mapped reads (mapq > 30) were retained and duplicate reads discarded. Contacts were then binned into matrices using Cooler (62). Analysis was performed on data pooled from two biological replicates, after confirming that the results of analysis of individual replicates were similar. Hi-C contact maps of OR clusters on chromosome 2 were generated with raw counts of Hi-C contacts normalized to counts/billion at 100kb resolution. The maximum value on the color scale was set to 150 contacts per 100kb bin. Analysis of zonal OR gene cluster contacts was performed using normalized counts binned at 50kb resolution. All analyses were repeated using balanced counts generated by Cooler (-mad-max 7).

### SXT sample preparation

Olfactory epithelium was microdissected into zones, dissociated using the Worthington Papain dissociation system, and sorted for OMP-GFP to create a single cell suspension of mature OSNs. Each batch for each zone was centrifuged and resuspended in 1x PBS. Single cells were prepared following the protocol described by Chen et al, 2022 (69). In brief, cells were loaded in 8-µm wide glass capillaries and cryopreserved after plunge-freezing in liquid nitrogen-cooled liquid propane. Samples were then stored in liquid nitrogen prior to data acquisition.

### SXT data collection

SXT was performed at the National Center for X-Ray Tomography (ncxt.org) on the beam line 2.1, located at Lawrence Berkeley National Laboratory, as described Chen et al., 2022 (69).

X-ray projection images were collected at an energy of 517 eV using a 50 nm resolution objective lens. During the data acquisition, samples were cryopreserved in a stream of liquid nitrogen-cooled helium. 92 projection images were sequentially captured at 2° increments around a 180° axis of rotation (capillary axis), with an exposure time of 300 ms. Cells located in areas of the capillary with diameter larger than 15 µm were collected using the half acquisition protocol, as described in Ekman et al., 2023 (70). 184 projection images were sequentially captured at 2° increments around a 360° axis of rotation, with an expo (71) sure time of 350 ms. Each tomogram was reconstructed using the algorithm AREC3D (71) and to calculate the linear absorption coefficient (LAC), each pixel intensity was normalized as described in Parkinson et al., 2012 (71).

### Tomogram segmentation

Segmentation of cellular structures was performed using semiautomated methods and visualized and refined in Amira 2021.2. software. The cell membrane was initially segmented using the algorithm ACSeg 3D (72) on the Biomedisa (https://biomedisa.info/) server (73). The nuclear envelope was manually segmented using the function “paintbrush” in Amira 2021.2.

### Partitioning Nucleus into Hetero-/Eu-chromatin

To segment the reconstructed linear absorption coefficient (LAC) of the nucleus into hetero and euchromatin, we used a smoothed Gaussian mixture modeling (GMM) approach. Using the manually segmented nucleus as a mask, a two-component GMM (74) was fit to the (mean) normalized LAC values. The model’s posterior probabilities were interpolated back into the volume and smoothed. The final segmentation was obtained by thresholding the smoothed probabilities at 0.5. Based on previous observation provided by Le Gros et al., 2016, mature olfactory neurons where selected base on the total amount of heterochromatin being above 37% of the total nuclear volume [47].

### Splitting Heterochromatin into Inner and Outer Compartments

Using a watershed approach, we segmented heterochromatin into inner and outer regions (75). First, a thin border region along the nuclear envelope (2 voxels thick) was used to extract the characteristic local thickness (76) of the peripheral heterochromatin. Seeds for the watershed segmentation were selected based on their distance to the nuclear envelope: voxels within the 50th percentile of the local thickness formed the outer seed, and voxels beyond the 90th percentile of the local thickness formed the inner seed. We removed small regions (less than 1% of the largest) from the inner seeds.

Using these seeds, we performed watershed segmentation on the Euclidean distance transform of the heterochromatin mask to partition the heterochromatin into distinct inner and outer compartments.

Segmentation of Heterochromatin Subregions by Recursive Thickness-Based Division: We developed a recursive watershed-based division method to partition connected inner heterochromatin regions into internally homogeneous subregions. Connected components were divided into separate regions if their connected throat was smaller than half of the mean thickness of the original cluster. This was done by initializing seeds from the EDT of the cluster and removing thin and small seeds (this was arbitrarily chosen as mean thickness one standard deviation above the cut threshold and a relative size larger than 1e-3 of the total volume) and applying watershed segmentation on the Euclidean distance transform of the original connected component. This step was repeated recursively on all separated clusters until convergence. The foci of each cell were determined to be the largest of these separated clusters. We inspected the result manually and corrected those that were wrong by selecting the cluster with the largest volume.

### Affinity Purification Mass Spectrometry

Cells were lysed in lysis buffer (150mM NaCl, 50mM Tris pH 7.5, 1% IGPAL-CA-630 Sigma #18896, 5% glycerol) on ice for 20 minutes, clarified, and total protein was quantified by BSA quantification. 5μg antibody (HP1a (HP1 alpha MA535397 ThermoFisher) and HP1b (D2F2, Cell Signaling) was added to 1μg total protein lysate per replicate. The cell lysates with antibody were incubated with Dynabead Protein A/G magnetic beads (Thermo Fisher Scientific) overnight at 4C. Supernatants were removed, beads were washed 2 times with wash buffer plus IGEPAL ca-630 (50 mM Tris pH 7.5, 150 mM NaCl, 5% glycerol, 0.05% IGEPAL), and twice with wash buffer (50 mM Tris pH 7.5, 150 mM NaCl, 5% glycerol). After the last wash, the beads were resuspended in 80 μL trypsin buffer (2 M Urea, 50 mM Tris pH 7.5, 1mM DTT, 5 μg/ml trypsin) to digest the bound proteins at 25 °C for 1 h at 1200 rpm. The supernatant was collected and the beads were washed twice with 60 μL Urea buffer (2 M Urea, 50 mM Tris pH 7.5) and the washes were combined with the supernatant. The combined elution was cleared of residual beads by a quick spin. 80 μL of the elution were used and disulfide bonds were reduced with 5 mM dithiothreitol (DTT), and cysteines were subsequently alkylated with 10 mM iodoacetamide. Samples were further digested by adding 0.5 μg sequencing grade modified trypsin (Promega) at 25°C. After 16 h of digestion, samples were acidified with 1% formic acid (final concentration). Tryptic peptides were desalted on C18 StageTips according to (77) and evaporated to dryness in a vacuum concentrator and reconstituted in 15 μL of 3% acetonitrile/ 2% formic acid for LC-MS/MS.

LC-MS/MS analysis was performed on a Q-Exactive HF. 5uL of total peptides were analyzed on a Waters M-Class UPLC using a 15cm Ion-Optics column (1.7um, C18, 75um x 15cm) coupled to a benchtop ThermoFisher Scientific Orbitrap Q Exactive HF mass spectrometer. Peptides were separated at a flow rate of 400 nL/min with a 90 min gradient, including sample loading and column equilibration times. Data were acquired in data-dependent mode. MS1 spectra were measured with a resolution of 120,000, an AGC target of 3e6 and a mass range from 300 to 1800 m/z. MS2 spectra were measured with a resolution of 15,000, an AGC target of 1e5 and a mass range from 200 to 2000 m/z. MS2 isolation windows of 1.6 m/z were measured with a normalized collision energy of 25.

Proteomics raw data was analyzed by MaxQuant v2.0.3.0 (78) using a UniProt database (Mus musculus, UP000000589), and MS/MS searches were performed under default settings with LFQ quantification. Data were further analyzed in R v3.6.3. Contaminants, and proteins only identified by site or reverse were removed, a pseudocount randomly taken from the bottom of the signal distributions was added to the LFQ intensity values and then the LFQ intensity values were log2 transformed. Proteins with a mean MS/MS count value for each IP condition below 5 were removed from subsequent analysis. Interacting proteins were identified as those that passed a log2FC sample IP over control IP cutoff of 1 and a p-value of 0.2. The p-value was estimated using a two-sided, two- sample Student’s t-test.

---

**Supplemental Figure 1.**
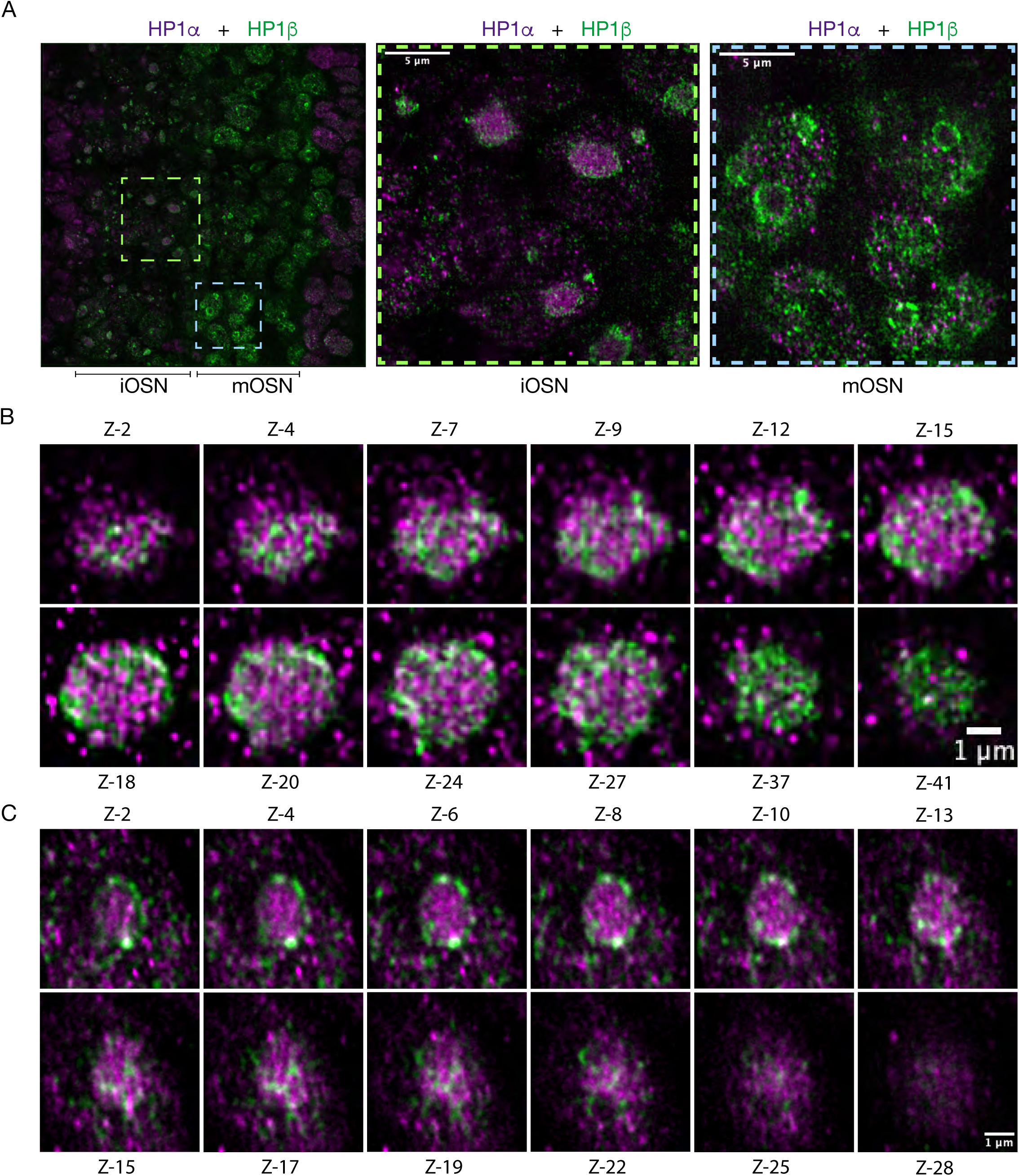
HP1α and HP1β segregate into different compartments. A) SoRA Spinning disk high resolution microscopy images. IF on MOE sections with HP1α (magenta) and HP1β (green) specific antibodies. Dashed line shows localization of zoom images on iOSN and mOSN cells. Scale bar is 5μm. B) and C) IF images from maximum projections from z stacks from an B) immature foci, or C) a mature foci. HP1α (magenta) and HP1β (green) occupy different compartments. Z-n indicates the z stack projection number. Scale bar is 1μm.

**Supplemental Figure 2.**
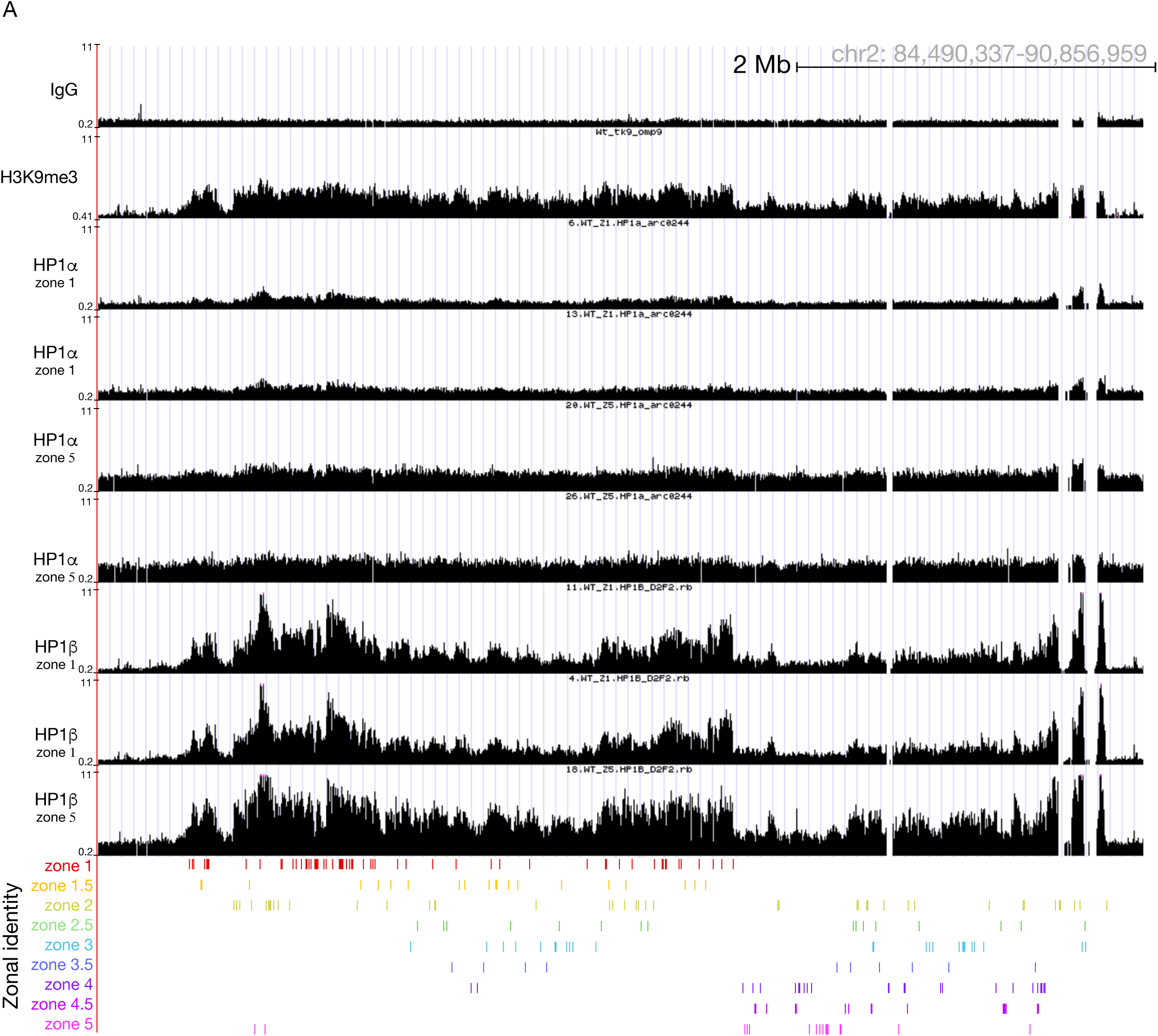
H**P**1β **is recruited to the OR clusters.** HP1α and HP1β ChIP-seq signal tracks on mature sorted OSN from micro dissected zones. Chromosome 2 OR cluster is shown as a representative image. Values are reads per 10 million. Below the signal tracks, OR genes are depicted in different colors indicating the assigned expression zone. Results are shown in duplicates and with the use of different antibodies except for HP1β zone5. Antibodies used on ChIP: H3K9me3 (Ab8898) HP1α (HP1 alpha MA535397 ThermoFisher) and HP1β (D2F2, Cell Signaling).

**Supplemental Figure 3.**
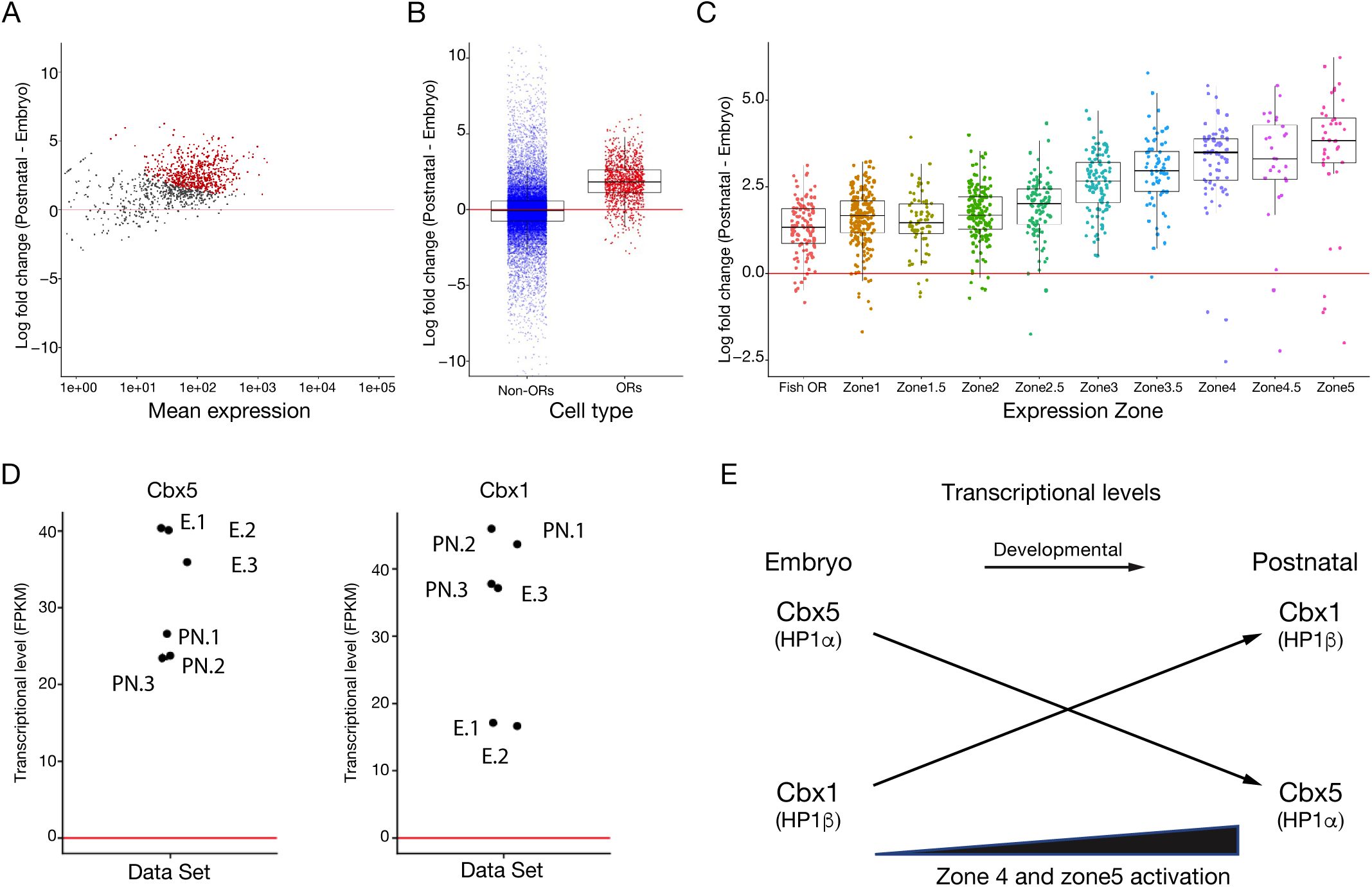
HP1α, HP1β and OR expression correlation analysis. A) RNA-seq analysis of gene expression of bulk RNA-seq in WT E17.5 embryos versus three week postnatal (PN) WT mice. Significantly changed genes are colored red (Padj < 0.05 for greater than 1.5-fold change, Wald test, n = 3). B) Box plot depicting gene expression of non-ORs and ORs in bulk RNA-seq (WT E17.5 embryos versus PN MOE). OR-gene activation increases in postnatal mice. C) OR gene expression from bulk RNA-seq reads bioinformatically separated into zones. Log2FC WT E17.5 embryos versus PN. D) Cbx1 and Cbx5 expression analysis shows a swap in the transcriptional levels on Cbx1 vs Cbx5 during development. E) Model summarizing our correlation analysis. In embryo, Cbx5 shows higher expression levels. In postnatal mice Cbx1 vs Cbx5 expression levels swap and receptors from ventral zones are expressed at higher levels.

**Supplemental Figure 4.**
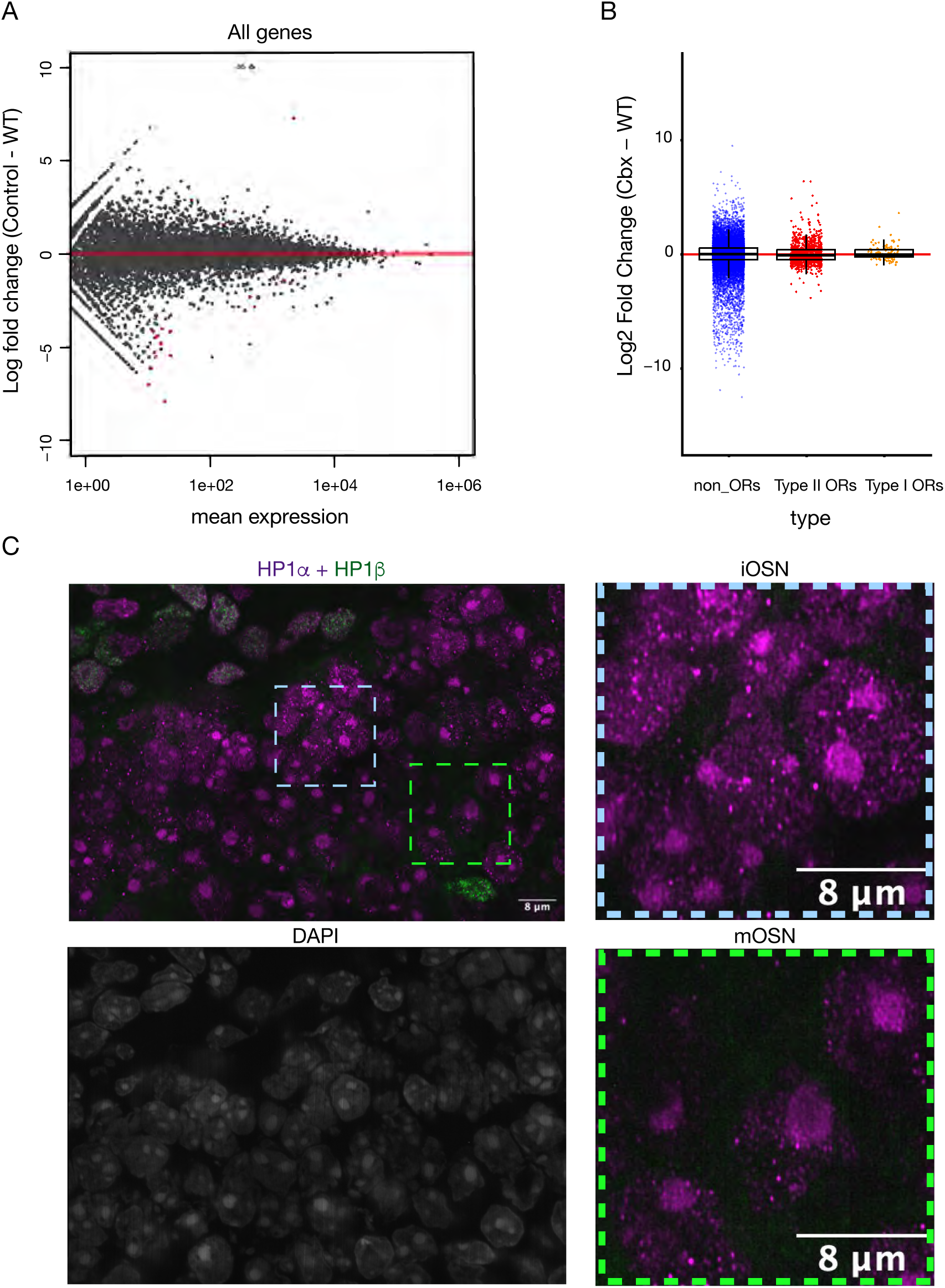
HP1α does not recapitulate HP1β staining. A) MA plot of RNA-Seq of Swap Knock-in Control mouse compared to WT mice. Significantly changed genes are colored red (Padj < 0.05 for greater than 1.5-fold change, Wald test, n = 3). B) Box plot of RNA-seq from knock-in mouse vs WT mouse showed no transcriptional or OR expression differences. C) SoRA Spinning disk high resolution microscopy images. IF on MOE sections with an HP1α (magenta) and HP1β (green) specific antibodies in zone-5 Swap mouse. Dashed line shows localization of zoom images on iOSN and mOSN cells. HP1α does not migrate to the periphery as HP1β does in mOSNs (futher analysis shown in figure 8). Antibodies used on IF: HP1α (ab109028) and HP1β (ab10811).

**Supplemental Figure 5.**
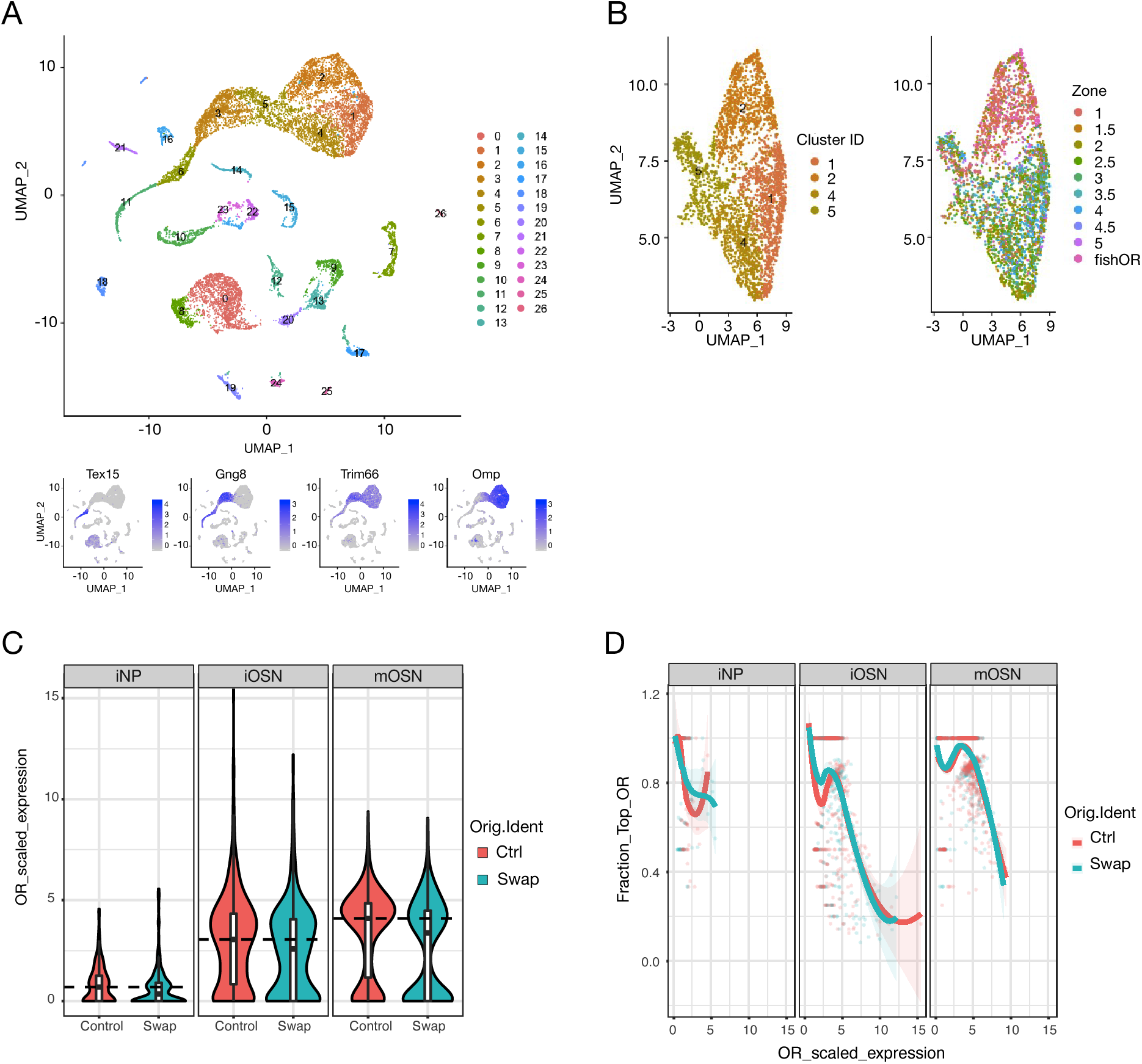
No effect on OR transcriptional levels and on the number of cells co-expressing ORs in Cbx Swap mice. A) Single Cell RNA-seq from total MOE segregated by Seurat cluster IDs. Bottom, expression levels of developmental markers. B) Left, mOSNs segregated by Seurat Cluster ID. Right, mOSNs segregated by zone. C) Violin plots depicting OR scaled expressed per cell in Control and Swap single cells, faceted by developmental stage. No differences in OR mRNA level per cell detected in the two genotypes. D) Fraction of OR counts from top OR, faceted by differentiation stage. No difference in number of cells co-expressing ORs was detected in the two genotypes.

**Supplemental Figure 6.**
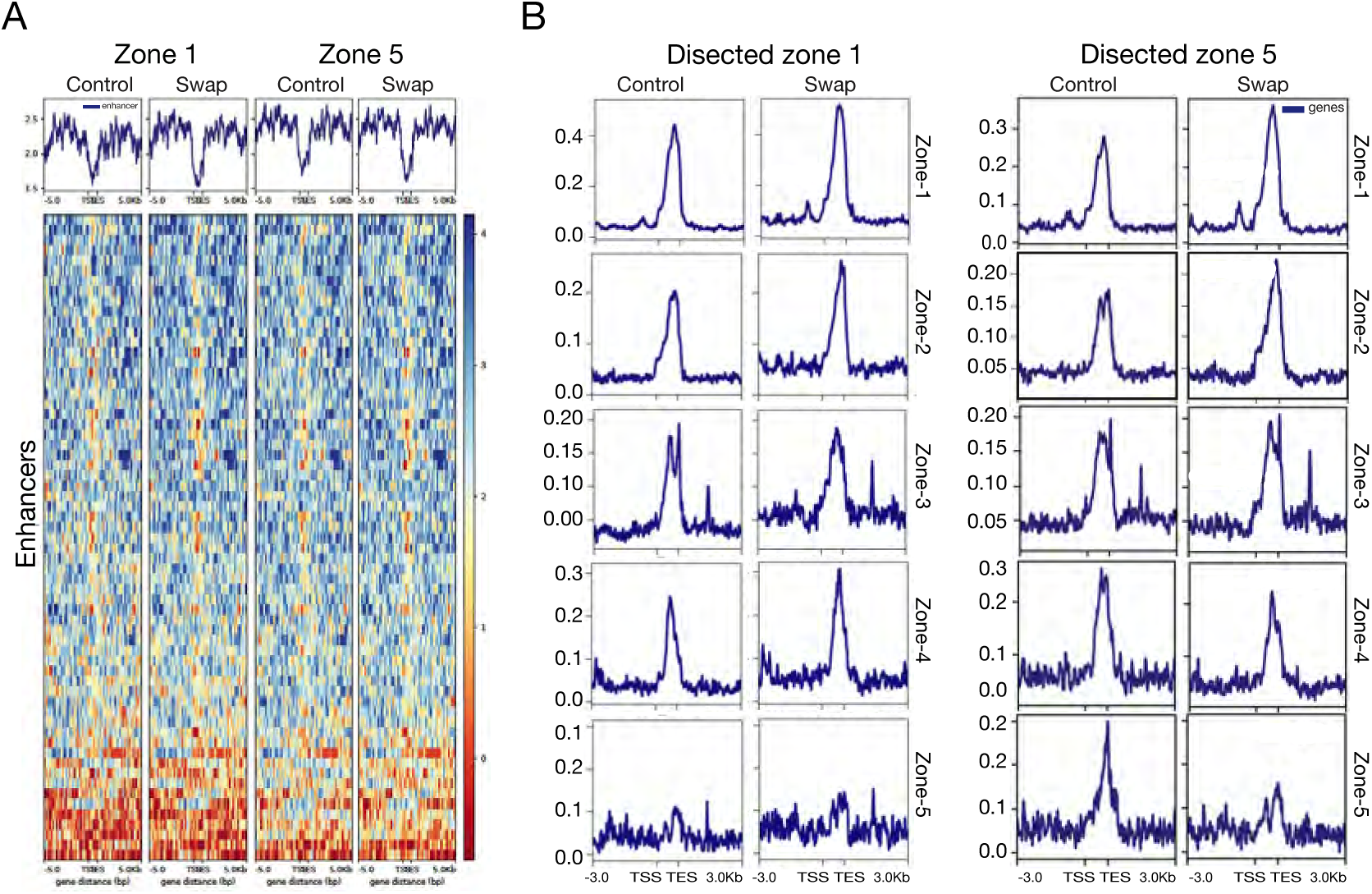
HP1β is implicated in H3K9me3 incorporation and MOE zone formation. A) Native ChIP-seq on micro dissected zone-1 vs zone-5 mature OSNs (n = 3 for each genotype), shows no changes in H3K9 methylation levels over Greek Islands. B) ATAC-seq on micro dissected zone-1 vs zone-5 showed a decrease in accessibility in repressed genes and an increase of accessibility in activated genes (n = 3 for each genotype).

**Supplemental Figure 7.**
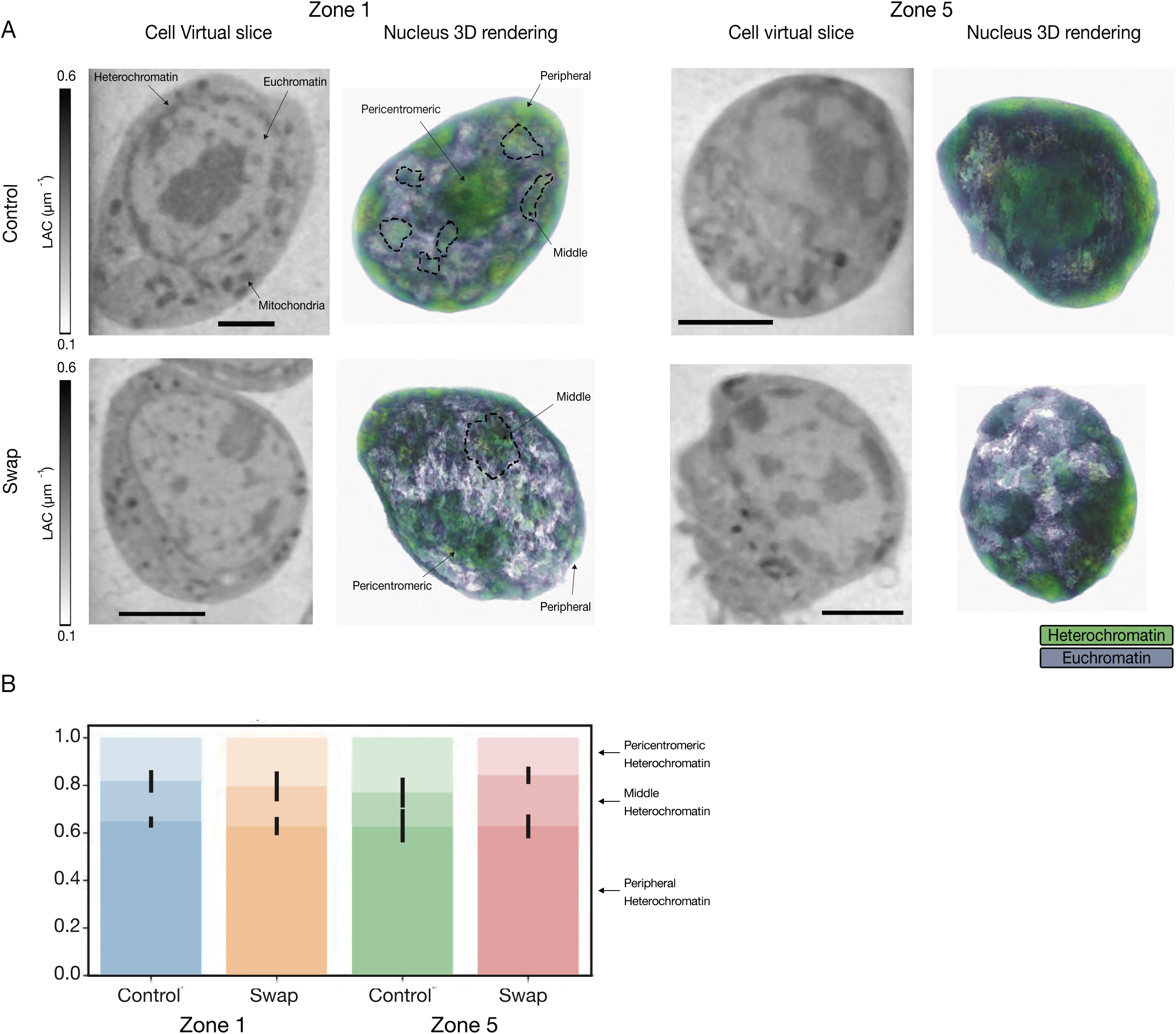
Effect of HP1β on chromatin architecture and nuclear organization. A) Cell virtual slice (left) and 3D rendering (right) of control (top) and swap mOSN cells (bottom) from micro dissected zone-1 and zone-5, imaged using soft X-ray tomography. The heterochromatin shows a typical LAC value of 0.28-0.02 μm-1 and is divided in three different areas: peripheral, pericentromeric and middle heterochromatin. The LAC values are reported in the range 0.1-0.6 μm-1 and the scale bar is 2 μm. B) Quantification of the heterochromatin distribution in the control and swap cells in zone 1 and zone 5. We report a trend towards increased middle heterochromatin in zone 5 swap cells as compared to the control cells in the same zone (p-value=0.066), with the control cells in zone 5 showing a larger volume of the pericentromeric heterochro-matin (p-value=0.061). No difference is observed between swap and control zone-1 cells.

**Supplemental Figure 8.**
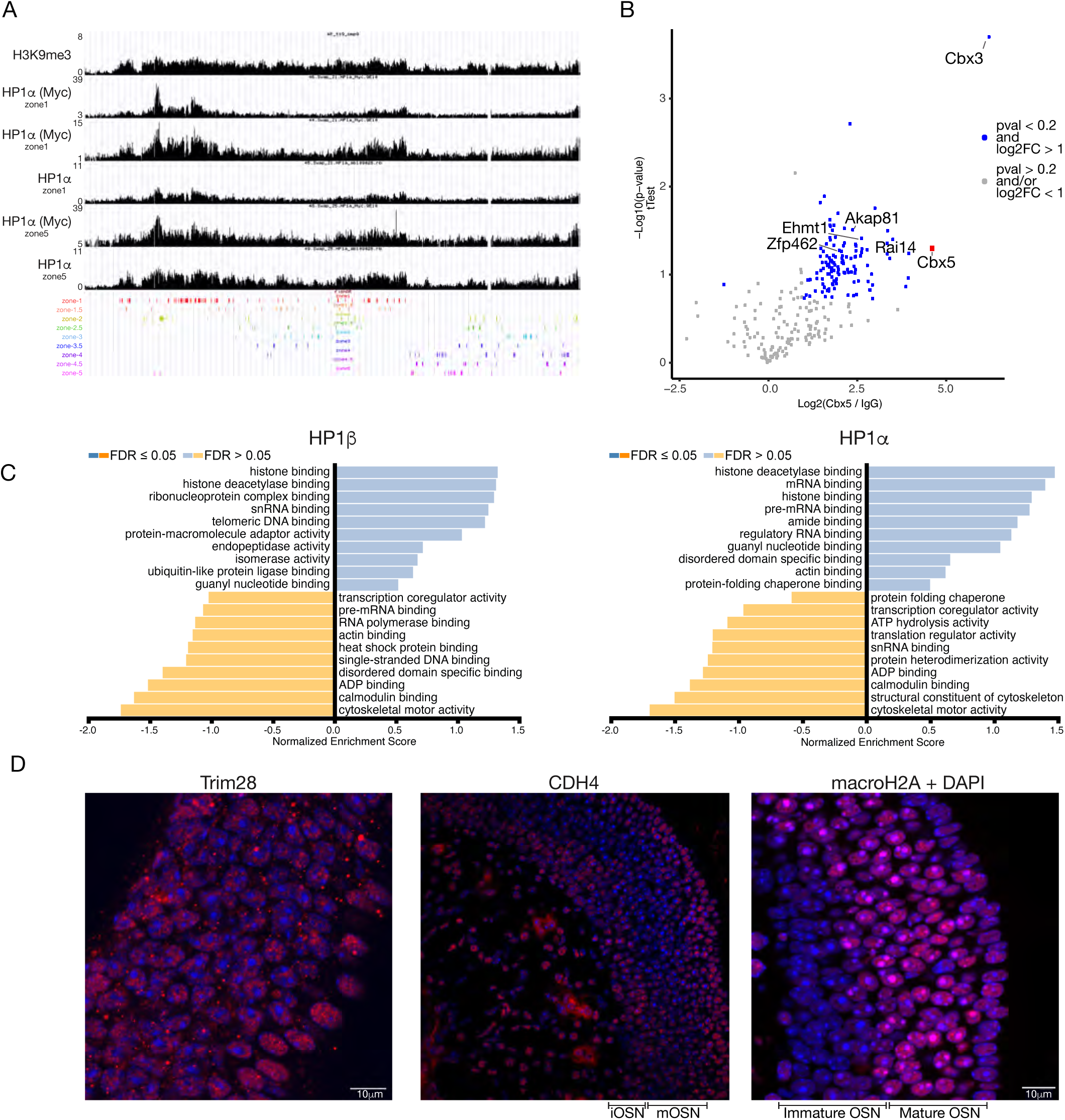
HP1α and HP1β interaction partners. A) ChIP-seq on sorted mOSN from micro dissected zone-1 vs zone-5. Signal tracks from representative example, chromosome 2 OR cluster is shown. Values are reads per 10 million. Below the signal tracks, OR genes are depicted in different colors indicating the assigned expression zone. Two different antibodies against HP1α and one against the Myc tag added to HP1α were used. B) Mass Spectrometry (MS) on samples from HP1α IP, using an specific antibody against HP1α. C) GSEA analysis on immuno-precipitated partners for HP1α and HP1β. FDR (False Discovery Rate). D) IF Imaging of MS candidates. Trim28 has a nuclear localization and localizes closer to the heterochromatin foci in mOSN. CDH4 has a nuclear localization and re-localizes with heterochromatin in mOSN. MacroH2A staining is reminiscent of HP1β in mOSN. Scale bar is 10μm. Antibodies used on IF: mH2A (ab208879), CDH4 (ab70469) and Trim28 (ab22553).

## Notes

### Competing Interest Statement

The authors have declared no competing interest.

